# The teosinte *mexicana* chromosomal inversion *Inv4m* modulates maize flowering time, plant height, and growth regulation gene networks

**DOI:** 10.64898/2026.06.16.732082

**Authors:** Fausto Rodríguez-Zapata, Hannah Pil, Nirwan Tandukar, Lauren Insko, Carolina Escalona-Weldt, Zehta Fazler, Alejandro Aragón-Raygoza, Melanie Perryman, Sergio Pérez-Limón, Sandra Senyo Fometu, Josh Strable, Daniel Runcie, Ruairidh Sawers, Rubén Rellán-Álvarez

## Abstract

Chromosomal inversions facilitate local adaptation by maintaining co-adapted allele complexes as a single inherited unit, but the genes driving their phenotypic effects are rarely identified. *Inv4m* is an inversion from the highland teosinte *Zea mays* ssp. *mexicana* that is nearly fixed at 2500 masl in Mexican traditional varieties but absent in temperate maize. While association studies link *Inv4m* to flowering time, its key adaptive trait, the mechanisms connecting it to this phenotype remain unknown. Here, we generate near isogenic lines by introgressing the highland Michoacán 21 (Mi21) *Inv4m* haplotype into temperate B73 through eight backcross generations. We assemble the *Inv4m*-Mi21 NIL genome and, aligning to three *Zea* reference genomes, provide the first precise delimitation of *Inv4m*, a 15.2 Mb region with breakpoints overlapping knob repeat arrays. In field trials across Pennsylvania and North Carolina, *Inv4m* consistently accelerated flowering, whereas its plant height effect reversed sign between regions, a genotype by environment interaction consistent with environment-dependent fitness. Using RNA-seq, we identify 465 differentially expressed genes, the strongest from a cluster of JUMONJI histone demethylases (JMJ). Comparing five *Zea* genomes, the B73 reference carries a lineage-specific tandem expansion of five JMJ paralogs while highland genotypes carrying *Inv4m* retain a single ancestral copy, a difference that accounts for most of the cluster’s differential expression. We then show *Inv4m* disrupts growth related coexpression modules, with the JMJ cluster losing connectivity, and traced a *trans* regulatory network linking it to cell proliferation genes including *pcna2* and *sec6*. In summary, we identify candidate genes and networks underlying *Inv4m*’s effects and propose that part of this inversion’s adaptive value may reside in a dosage-sensitive regulator whose action propagates through a downstream network of genes that are involved in growth and flowering and are important for highland adaptation. Recombination suppression may thus protect a co-adapted regulatory architecture rather than independent alleles alone.

**Graphical Abstract:** 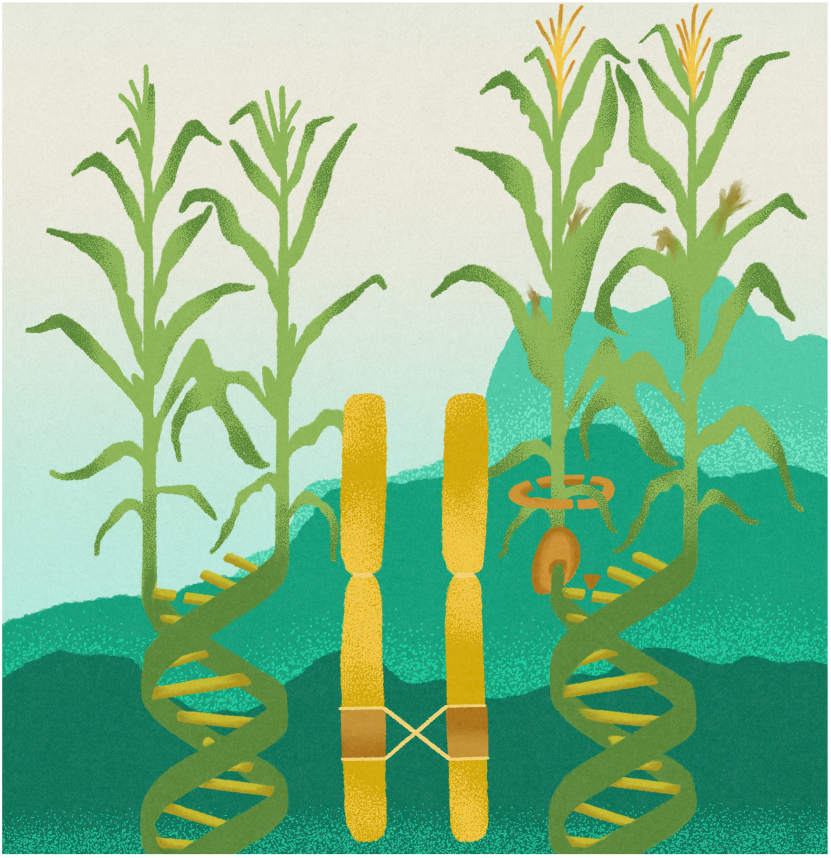

## Introduction

CHROMOSOMAL inversions are a widespread form of structural variation, segregating in natural populations of plants, animals, fungi, and humans (Wellenreuther and Bernatchez 2018a; Kirkpatrick and Barton 2006; Kirkpatrick 2010). By suppressing recombination between standard and inverted haplotypes, inversions can maintain coadapted allele complexes as single inherited units, a mechanism originally proposed by Dobzhansky (1971) and now central to theories of local adaptation, supergene maintenance, and incipient speciation (Kirkpatrick and Barton 2006; Kirkpatrick 2010; Lowry and Willis 2010). A growing body of work establishes that megabase scale inversions are repeatedly associated with environmental clines and adaptive phenotypes across diverse taxa. Yet despite their evolutionary prominence, the genes within inversions that actually drive their phenotypic effects remain unidentified in most systems: the same recombination suppression that allows inversions to maintain coadapted complexes also confounds genetic dissection of the loci they contain.

Maize (*Zea mays* ssp. *mays*) and its wild relatives are a powerful system for tackling this problem. Around 9,000 years ago, indigenous groups in the tropical lowlands of the Balsas River basin in Mexico domesticated maize from its wild relative, teosinte *parviglumis* (*Zea mays* ssp. *parviglumis*) (Matsuoka *et al*. 2002; Piperno and Flannery 2001). Recently domesticated maize was introduced to the Mexican highlands, bringing it into sympatry with the wild highland teosinte *mexicana* (*Zea mays* ssp. *mexicana*) and facilitating the introgression of highland adaptive alleles (Hufford *et al*. 2013; Calfee *et al*. 2021; Barnes *et al*. 2022), which preadapted maize for its subsequent expansion into temperate zones (Yang *et al*. 2023). However, while most modern maize contains significant introgression of teosinte *mexicana* (Yang *et al*. 2023), not all highland adaptive loci are present in temperate maize. Highland-associated chromosomal inversions such as *Inv4m* and *Inv9f* are prevalent in highland teosinte populations (Pyhajarvi *et al*. 2013) and traditional Mexican highland maize varieties (Crow *et al*. 2020; Gonzalez-Segovia *et al*. 2019) but are rare in temperate maize.

*Inv4m* is the best characterized highland adaptive inversion in maize. A previous analysis of teosinte populations using the Maize 50K chip inferred that *Inv4m* spans 13 Mb, is predominantly found in teosinte *mexicana* populations (Pyhajarvi *et al*. 2013), and is introgressed into sympatric highland maize populations (Pyhajarvi *et al*. 2013; Calfee *et al*. 2021; Hufford *et al*. 2013) In Mexican traditional varieties, *Inv4m* is associated with early flowering time (Romero Navarro *et al*. 2017), reduced genetic diversity, reduced recombination (Gonzalez-Segovia *et al*. 2019) and is nearly fixed at 2500 masl (Crow *et al*. 2020). Mexican highlands have phosphorus deficient, low pH volcanic soils, and traditional highland varieties show high phosphorus use efficiency (Bayuelo-Jiménez and Ochoa-Cadavid 2014). Because *Inv4m* carries a key phosphorus transporter (Salazar-Vidal *et al*. 2016) and is fixed in highland maize, its adaptive role had long been hypothesized to involve improved performance in low phosphorus soils, a hypothesis we reject in a companion study (Rodríguez-Zapata *et al*. 2026). Crucially, *Inv4m* displays the hallmark of local adaptation through antagonistic selection: plants carrying the highland allele flower earlier and yield more above 2000 masl, while the same allele delays flowering and reduces yield below 1000 masl (Crow *et al*. 2020).

Despite strong evidence linking *Inv4m* to local adaptation, the physiological processes and environmental factors underlying its adaptive role and the genetic mechanisms by which genes within the inversion drive its adaptive phenotypic effect have remained unclear. Population genetic scans, association studies, and growth chamber expression analyses have implicated candidate regions but have not identified the molecular mechanisms underlying adaptive traits. Highland *Inv4m* upregulates photosynthesis genes in response to cold at the seedling stage (Crow *et al*. 2020) and is associated with earlier flowering in the Mexican highlands, which likely enhances fitness in environments with limited growth degree accumulation throughout the year (Romero Navarro *et al*. 2017; Eagles and Lothrop 1994; Giauffret *et al*. 2000). Crow *et al*. (2020) identified differentially expressed genes between *Inv4m* haplotypes in cold stressed seedlings under growth chamber conditions, with transcriptomic effects enriched for photosynthesis and lipid metabolism, but the field-relevant phenotypic consequences of these expression changes, and how they propagate through gene regulatory networks, have not been established. More broadly, the effects of *Inv4m* introgression on gene expression and phenotype in temperate field environments have not been directly tested.

Here we close this gap with a multi level characterization of *Inv4m* that combines careful control of genetic background, phenotyping across multiple common gardens to capture genotype by environment effects, and detailed expression analysis to dissect regulatory architecture. We introgressed the highland inverted haplotype from the Mexican traditional variety Mi21 (Hayano-Kanashiro *et al*. 2009) into B73 (Schnable *et al*. 2009), a temperate maize inbred line, through eight generations of backcrossing to generate BC_8_S_2_ near isogenic lines (NILs). We then used long read assemblies of highland materials and the Mi21 NIL to precisely delimit *Inv4m* breakpoints. We found that *Inv4m* breakpoints overlap chromosomal knob repeat arrays consistent with an ectopic recombination origin. *Inv4m* modulated flowering time and plant height across our field environments and showed genotype by environment interactions and produced elongated shoot apical meristems in seedlings, a developmental signature of accelerated flowering (Danilevskaya *et al*. 2008; Wu *et al*. 2008; Leiboff *et al*. 2015; Thompson *et al*. 2015). We then traced the most significant gene expression difference to a cluster of JUMONJI histone demethylases (JMJ) within *Inv4m* that show copy number variation between highland and temperate materials. Coexpression analysis revealed that *Inv4m* introgression rewires native B73 networks: the *jmj2*/*jmj4* cluster loses hub connectivity in growth related coexpression modules, and a *trans*-regulatory network connects the cluster to cell proliferation genes including *pcna2* and *sec6*. This points to a working hypothesis in which JMJ dosage modulates growth and flowering time through chromatin modification of target genes altering their expression patterns.

Inversions are classically thought to become adaptive either by capturing epistatically interacting alleles (Dobzhansky 1971) or by sheltering locally favored allele combinations from gene flow (Kirkpatrick and Barton 2006; Kirkpatrick 2010). *Inv4m*’s elevation-dependent fitness reversal (Crow *et al*. 2020) fits the local adaptation framework; recombination suppression precludes the dissection of epistasis required to test the supergene hypothesis. Our results point to a mechanism both theories implicitly assume: some of *Inv4m*’s effects may flow through a single dosage-sensitive regulator whose action propagates through a downstream gene network that can extend beyond the inversion’s genomic limits. Recombination suppression would then protect a regulatory architecture in *Inv4m* connecting a series of genes involved in growth and flowering time, likely via the action of JMJ histone demethylases and not simply a collection of independent alleles. While our results point to molecular mechanisms underlying *Inv4m*’s mode of action, fully dissecting these mechanisms will require further molecular characterization and ultimately restoration of colinear haplotypes using genomic engineering approaches (Schwartz *et al*. 2020) for fine mapping within the inversion.

## Results

### Inv4m is a 15 Mb inversion delimited by breakpoint segments overlapping chromosomal knob repeats

We assembled the Mi21 NIL genome from PacBio HiFi reads (*∼*14x coverage) using hifiasm (Cheng *et al*. 2021a) and scaffolded contigs against B73 and PT references with RagTag (Alonge *et al*. 2022) (N50 = 13.7 Mb; 2.2 Gb total). Chromosome 4 was represented by a 229 Mb primary scaffold covering B73 chr4 positions 18–250 Mb, with the short arm tip (0.6–17 Mb) on a separate 17.2 Mb scaffold, a split confirmed by AnchorWave (Song *et al*. 2022) alignments to both B73 and PT. Gene annotations were transferred from B73 via Liftoff (Shumate and Salzberg 2021).

To delimit the recombination breakpoints of *Inv4m*, we performed CDS-anchored whole genome alignments with AnchorWave (Song *et al*. 2022) on chromosome 4. We aligned the Mi21 NIL assembly, PT, a close relative to the *Inv4m* donor parent Mi21, representing the Mexican highlands (Perez-Limón *et al*. 2022; Vielle-Calzada *et al*. 2009), and TIL18 (highland teosinte *mexicana*) to B73, the recurrent parent used for *Inv4m* introgression, (Fig 1B; Supplementary Table S2). In each pairwise comparison, the *Inv4m* region aligned to the B73 minus strand while flanking regions aligned to the plus strand, as expected for an inversion (Fig 1C). This allowed us to narrow down *Inv4m* to between genes *Zm00001eb190470* and *Zm00001eb194800*, spanning 15.2 Mb and 432 annotated genes in the B73 v5 genome. The same bounding genes delimited a 13.4 Mb inversion in PT, a 13.2 Mb inversion in TIL18, and a 10.9 Mb inverted segment in the Mi21 NIL (Supplementary Table S2; Fig 1C). The inversion was not bounded by collinear (plus/plus) alignments on either side. Instead, each breakpoint coincided with an unaligned segment, i.e. a gap in the AnchorWave alignment where no CDS anchors could be placed. These breakpoint segments were largely devoid of annotated genes; only *Zm00001eb190490* was present, in the upstream segment of B73. To characterize the sequence composition of these gene poor breakpoint segments, we generated self-versus-self dotplots using LAST (Kiełbasa *et al*. 2011). The dotplots showed prominent tandem repeat arrays at both breakpoints in all four genomes (Fig 1D), prompting us to examine the available repeat annotation of B73. The repeat annotation available in MaizeGDB for B73 identified the tandem arrays at the breakpoints as chromosomal knob associated repeats of the *knob180* and *TR-1* classes. The dotplots, however, showed tandem repeat signal extending beyond the annotated repeat boundaries, suggesting that more divergent copies were not captured by the existing annotation (Fig 1D). We therefore retrieved the original *knob180* (Dennis and Peacock 1984) and *TR-1* (Hsu *et al*. 2003) reference sequences and performed BLAST+ (Camacho *et al*. 2009) megablast searches against each chromosome 4.

**Figure 1.**
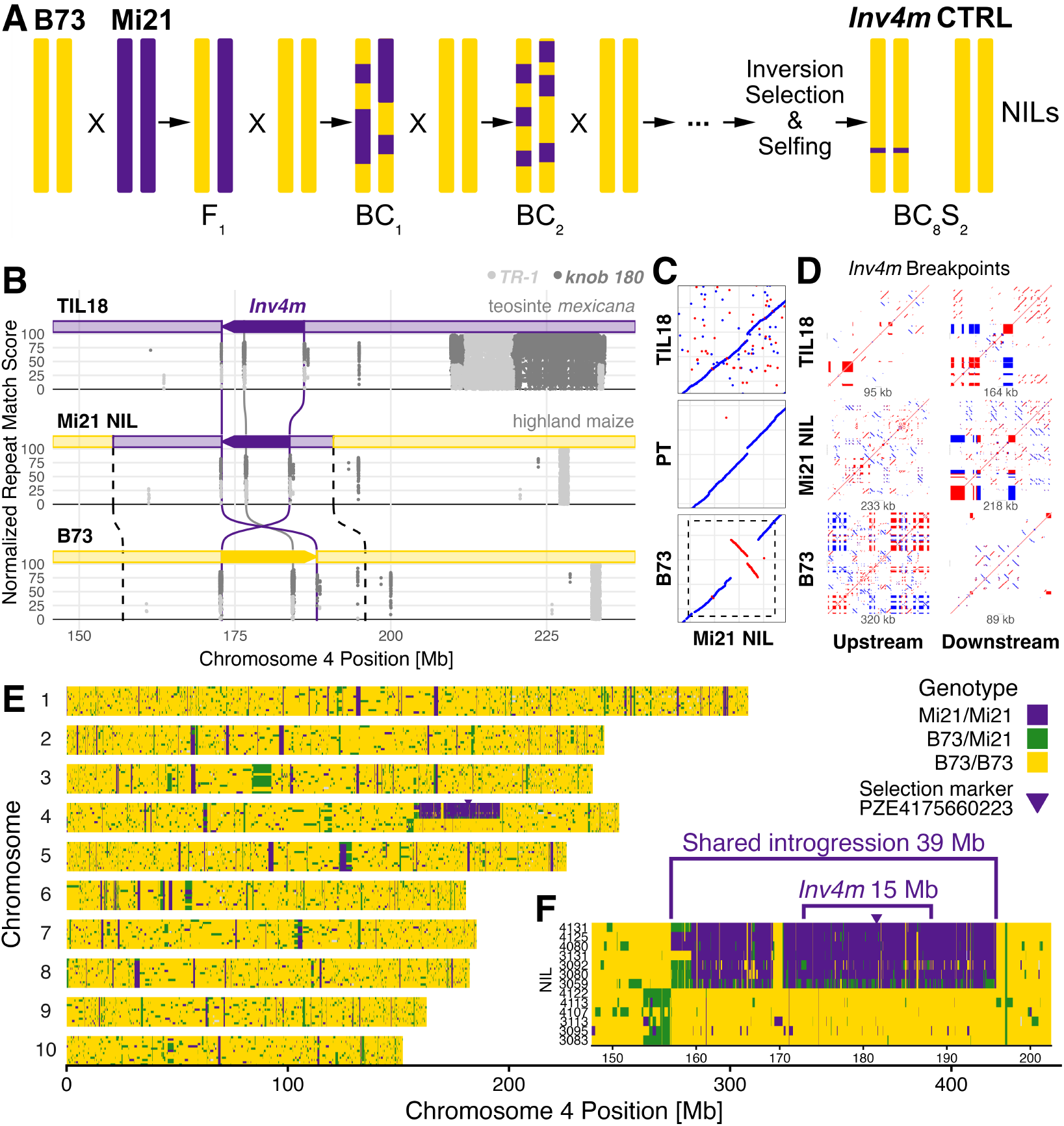
Chromosomal Inversion *Inv4m*: Delimitation, Introgression Breeding Design, Field Experimental Setup and Population Genotype. **(A)** Near Isogenic Line population breeding scheme. **(B)** Distribution of 180 bp knob repeats and *TR-1* elements across the *Inv4m* region. Each point represents a BLAST+ (Camacho *et al*. 2009) hit (normalized match score). Solid purple lines mark inversion breakpoints; dashed black lines mark the estimated *mexicana* introgression boundaries in Mi21 NIL and B73, connected by sigmoid curves reflecting the coordinate transformation between genomes. **(C)** AnchorWave (Song *et al*. 2022) whole genome alignment dotplots of Mi21 NIL chromosome 4 versus TIL18 (teosinte *mexicana*), PT (highland maize), and B73 (reference). Collinear anchors in blue, inverted in red. The dashed rectangle on the B73 panel marks the estimated introgression extent (*∼*141– 170 Mb in Mi21 NIL; *∼*157–195.9 Mb in B73 coordinates), determined from positional offset analysis of the alignment. **(D)** Sequence self-similarity dotplots, using LAST (Kiełbasa *et al*. 2011), for the inversion breakpoint segments as defined in Supplementary Table S2. Forward matches in red, reverse in blue. Rectangles indicate tandem repeats. **(E)** NIL genotypes at BC_8_S_2_, of plants used for RNA analysis, *n* = 13, B73 NAM5 coordinates. Genotype color: gold = B73/B73, green = B73/Mi21 (heterozygous), purple = Mi21/Mi21. **(F)** Zoom into *Inv4m* introgression. Plants are sorted by genotype at the *Inv4m* tagging SNP PZE04175660223 (181.6 Mb, downwards pointing triangle) then by field row number.

This search detected knob repeats in the breakpoint segments, inside the inversion, and elsewhere on chromosome 4 in all four genomes (Fig 1B). Knob repeats were the most abundant repeat class within the breakpoint segments (Supplementary Table S3).

### Divergent highland introgression is mostly confined to a 39 Mb segment spanning Inv4m

We generated BC_8_S_2_ NILs by backcrossing Mi21 into B73 for eight generations and selfing, selecting at each cycle for the highland allele at an inversion tagging CAPS marker (Methods) (Fig 1A). RNA-Seq-based genotyping of 13 BC_8_S_2_ individuals and identical genotype run length analysis (Layer *et al*. 2016) defined a 39 Mb shared introgressed segment around *Inv4m* (B73 v5 chr4: 157.0–195.9 Mb; Fig 1E, F), of which 15 Mb corresponds to the inversion itself and 24 Mb to flanking linkage drag. Inversion-free lines were obtained by counter selecting for the B73 allele at the same marker and used as controls. The 24 Mb of flanking drag is a substantial reduction from the BC_5_ populations of Crow *et al*. (2020), where residual donor sequence extended over 57 Mb. Detailed SNP distributions, per-plant introgression statistics are provided in Supplementary Fig S1.

### Inv4m introgression modulates flowering time and plant height

At the Pennsylvania 2022 site (PA2022), highland *Inv4m* plants flowered *∼*1.3 days earlier and were*∼* 6.5 cm taller than control lines, with a modest increase in harvest index and additional significant effects on stover dry weight and cob diameter (Fig 2A). We then crossed *Inv4m* Mi21 NILs with the inbred W22 to test the effect of *Inv4m* in hybrids. *Inv4m* hybrids were taller and had higher yield components such as total grain weight (Fig 2B). To test possible G x E effects across environments, we phenotyped the same NILs at two additional location-years (PA2025) in the same PA location as the previous one and North Carolina (NC2025). Two-way ANOVA detected significant genotype*×* environment interactions for all phenotypes (Supplementary Table S4; Supplementary Fig S3), strongest for plant height. Flowering acceleration was consistent across all three environments. In contrast, the height effect reversed sign: Cohen’s *d* ranged from +1.45 to +2.23 in the two Pennsylvania environments but was *−*1.28 in North Carolina (Supplementary Fig S4).

**Figure 2.**
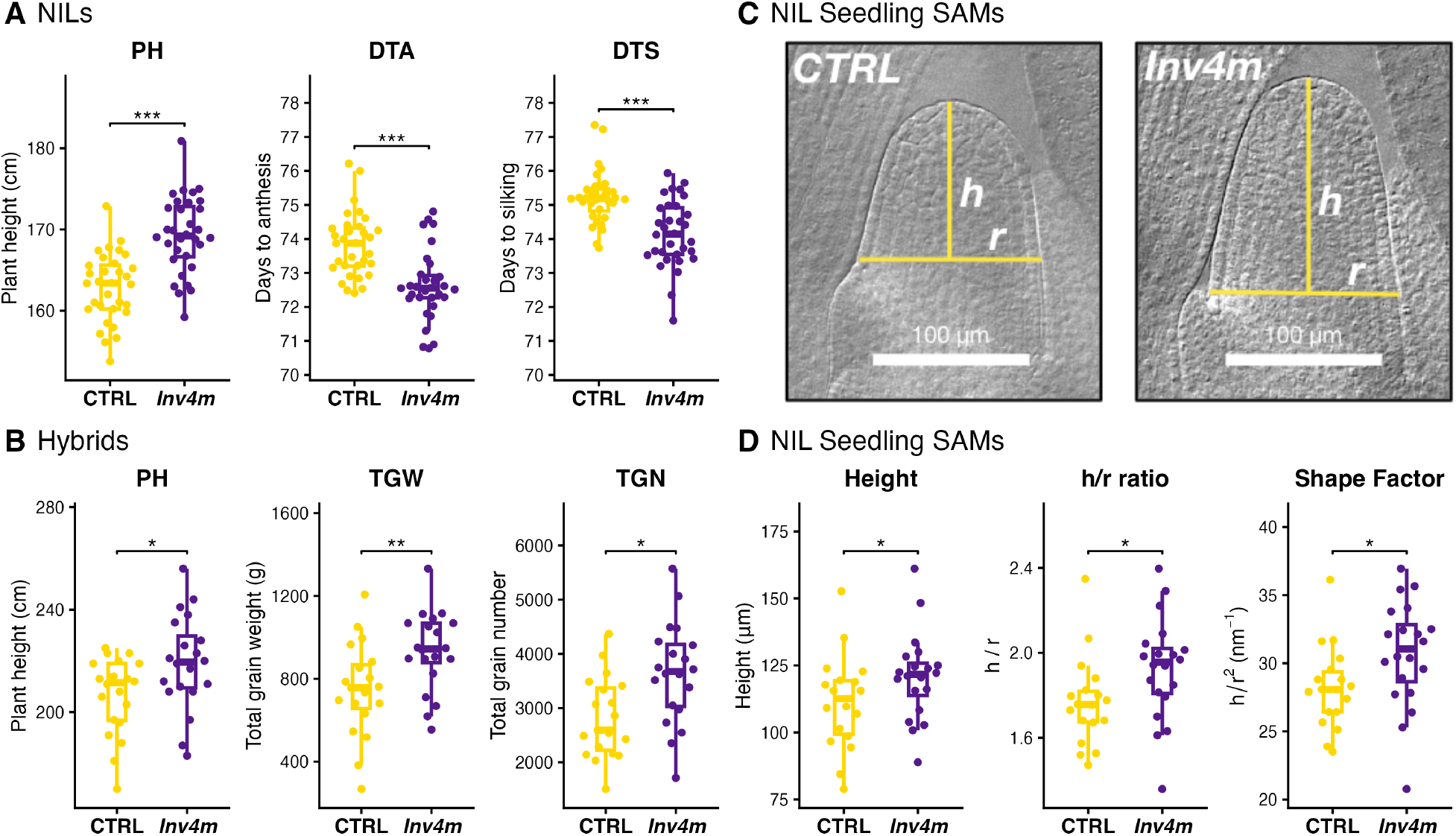
Effect of *Inv4m* on field phenotypes and shoot apical meristem architecture. CTRL (yellow) vs *Inv4m* (purple). **(A)** PA2022 NIL field trial; *t*-tests for marginal genotype contrasts. **(B)** PA2024 W22 x NIL hybrid field trial; *t*-tests for contrasts between hybrids with either *Inv4m* or CTRL NIL parents. **(C)** Differential interference contrast micrographs of cleared shoot apical meristems from seedlings grown two weeks in a growth chamber; yellow lines mark height (*h*) and radius (*r*). Scale bars: 100 *µ*m. **(D)** SAM dimensions: height, height/radius ratio, and shape factor; one tailed Welch *t*-tests (*Inv4m >* CTRL). Significance: FDR adjusted ^*\**^ *p <* 0.05, ^*\**^ *p <* 0.01, ^*\*\**^ *p <* 0.001, ^*\*\**^ *p <* 0.0001.

To characterize the architectural basis of this reversal, we measured internodes in NC2025, where *Inv4m* plants were shorter. Mean internode length was similar between genotypes, but the difference was concentrated at positions 2–4 from the top, with position 2 (immediately below the tassel) accounting for the majority of the height difference (Supplementary Fig S5). Aboveground node count did not differ between genotypes, indicating that *Inv4m* affects plant height through differential elongation of specific internodes rather than through changes in phytomer number.

To test whether the *Inv4m* flowering time effect generalizes across donor genetic backgrounds, we analyzed an independent B73 x Teosinte BC2S3 NIL panel (Zea Exotic Allele Library) grown at the same North Carolina site (NC2025) in which the *Inv4m* bearing segment had been introgressed into B73 from multiple teosinte donor accessions spanning four *Z. mays* ssp. *mexicana* ancestry groups (Chalco, Durango, Mesa-Central, Nobogame) and *Z. mays* ssp. *huehuetenanguensis* (Huehuetenango). A linear mixed model with strictly nested random effects (ancestry / BC2 family / NIL; see Methods) replicated the flowering time signal we observed in the Mi21 single donor panel: plants carrying the donor *Inv4m* inverted karyotype silked 0.88 ± 0.31 days earlier and reached anthesis 0.61 *±* 0.27 days earlier than their recurrent (B73 standard karyotype) siblings within the same NIL line (Supplementary Fig S2A,B). The contrast was driven primarily by the Durango ancestry, which provided both the largest donor sample (n = 37 plots) and the largest per ancestry effect (DTS: *−*1.47 *±* 0.46 d, FDR = 0.009); the *Inv4m×* ancestry interaction was not significant for either trait (*p >* 0.2), so the data do not support claiming the effect varies meaningfully by donor ancestry. Plant height showed a positive but non significant donor effect in the ZeaL panel (+1.6 *±* 2.7 cm, *p* = 0.55; Supplementary Fig S2C,D).

To investigate whether these whole plant phenotypes were associated with changes in meristem organization, we examined shoot apical meristems (SAMs) from two week old seedlings grown in controlled environment chambers (Fig 2C,D). *Inv4m* seedlings showed a 9% increase in SAM height with width unchanged. Elevated shape metrics (*h*/*r* and *h*/*r*^2^) confirmed the elongation, a developmental signature consistent with the larger, more elongated meristems previously associated with accelerated flowering (Danilevskaya *et al*. 2008; Wu *et al*. 2008; Leiboff *et al*. 2015; Thompson *et al*. 2015).

### Leaf developmental stage drives global transcriptional changes while Inv4m effects are predominantly restricted to the Inv4m region

To characterize the transcriptional landscape of *Inv4m* introgression, we performed RNA-seq on leaves sampled across a developmental gradient from field grown NILs (Fig 3A). Multidimensional scaling (MDS) of gene expression identified three major axes of variation (Fig 3B). The first two dimensions captured 39% of total variance, corresponding to phosphorus treatment (27%) and leaf developmental stage (12%), as described in detail in our companion study (Rodríguez-Zapata *et al*. 2026). The third dimension explained 8% of variance and strongly correlated with genotype (*r* = 0.92, *FDR* = 2.06*×* 10^*−*17^), clearly separating *Inv4m* from CTRL samples.

**Figure 3.**
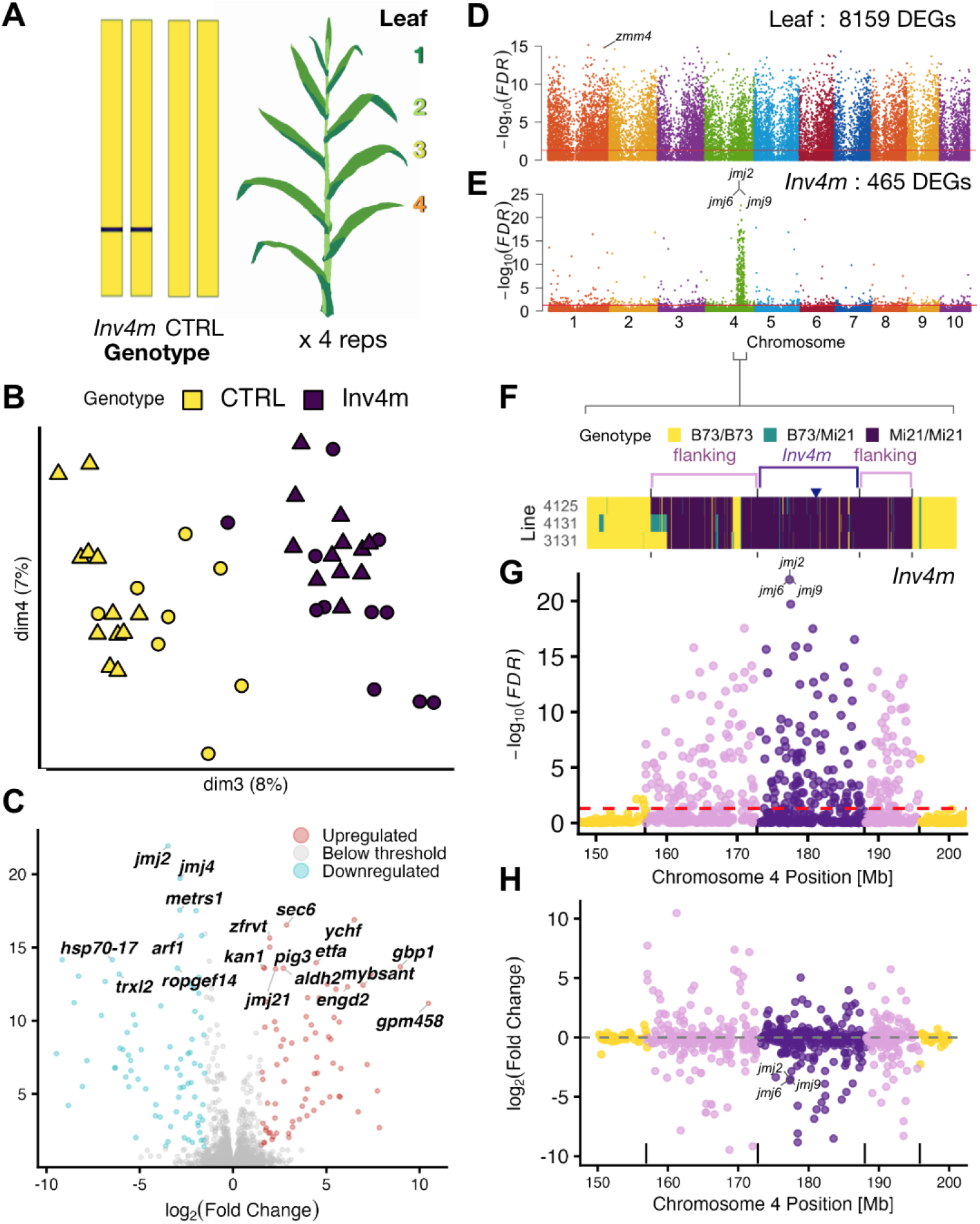
Global and local transcriptomic effects of *Inv4m* across leaf stages. **A)** Leaf sampling strategy. Four developmental leaf stages (1 to 4) were sampled across four replicates for both *Inv4m* and CTRL genotypes. **(B)** Multidimensional Scaling Analysis of the leaf transcriptome. Samples are colored by genotype (CTRL: yellow; *Inv4m*: purple) and shaped by leaf stage, showing clear separation along dimension 3. **(C)** Volcano plot of *Inv4m* associated DEGs. Labeled genes highlight significant perturbations, including the downregulation of multiple JUMONJI histone demethylases within the inversion. **(D)** Global Manhattan plot showing Differentially Expressed Genes (DEGs) associated with leaf stage across the ten maize chromosomes. **(E)** Global Manhattan plot for DEGs associated with the *Inv4m* genotype effect. **(F)** Haplotype map of the introgressed region on chromosome 4 for representative NIL lines. The blue triangle indicates the tagging SNP PZE04175660223 used for selection. **(G)** Zoom into the 150-200 Mb region of chromosome 4, showing the effect of *Inv4m* introgression in gene expression. **(H)** Expression effect of *Inv4m* across chromosome 4. Moving average lines green for upregulated, red for downregulated.

The spatial distribution of differentially expressed genes (DEGs) showed a striking contrast between leaf stage and *Inv4m* effects (Fig 3C-E). Leaf developmental stage produced a genome wide transcriptional response, with significant DEGs distributed across all ten chromosomes (Fig 3D). In contrast, the *Inv4m* effect was sharply localized: a prominent peak of differential expression on chromosome 4 corresponded precisely to the introgressed region (Fig 3E-F). We classified the 465 *Inv4m* responsive DEGs (*FDR <* 0.05) by their position relative to the inversion (Supplementary Table S1). DEGs within the shared introgression (39 Mb spanning *Inv4m* and flanking regions) showed 97-fold enrichment compared to the rest of the genome (Fisher’s exact test, *p <* 10^*−*200^). This enrichment was even larger among strong DEGs (| log_2_ FC| *>* 1.5): the 116 strong DEGs in the shared introgression represented a 141-fold enrichment over genomic background. DEG density did not differ between *Inv4m* proper and the flanking introgressed regions (odds ratio = 1.02, *p* = 0.95), indicating that the transcriptional perturbation extends throughout the introgressed segment rather than concentrating at the inversion, or the flanking regions.

To test whether differential expression in the introgression is explained by coding sequence divergence between the B73 and Mi21 haplotypes, we computed CDS percent identity from LAST pairwise alignments for 254 expressed genes with ortholog hits in the shared introgression. Genes inside the inversion had significantly higher CDS divergence than flanking genes (median 1.1% vs 0.75%; Wilcoxon *p* = 0.0034; Supplementary Fig S7A), consistent with suppressed recombination preserving ancestral *mexicana* divergence within the inversion. An additive linear model (|log_2_FC| *∼* CDS divergence + region) explained 10.4% of the variance in expression change (adj. *R*^2^ = 0.104; divergence *p* = 2.4e-06; region *p* = 0.0023; Supplementary Fig S7B). After accounting for divergence, genes inside the inversion showed smaller absolute expression changes than flanking genes, indicating that the inversion dampens rather than amplifies expression divergence. Notably, 21 genes had identical CDS between B73 and Mi21, and all were significantly differentially expressed (FDR *<* 0.05). Of these, 20 of 21 were located in the flanking region, a significant overrepresentation (Fisher’s exact test, *p* = 0.00013, OR = 0.1). Because these genes lack coding sequence divergence, their differential expression is most parsimoniously attributed to *trans*-regulatory effects originating from divergent loci elsewhere in the introgression.

The peak of differential expression within *Inv4m* proper corresponds to a cluster of *JUMONJI* histone demethylases comprising *jmj2, jmj6*, and *jmj4* (Fig 3G-H). The NAM v5 annotation consolidates *jmj2, jmj6*, and *jmj9* as alternative transcripts of a single locus (*Zm00001eb191790*), while *jmj4* is annotated separately (*Zm00001eb191820*). These genes showed the strongest significance in *Inv4m* plants, with *jmj2* exhibiting a log_2_ FC = *−*3.49 (*FDR <* 10^*−*20^; Supplementary Table S6).

### Inv4m introgression disrupts extant coexpression modules of the B73 genomic background

To assess whether *Inv4m* causes network level perturbations beyond individual DEGs, we performed weighted gene coexpression network analysis (WGCNA) (Shahan *et al*. 2018) on the 465 DEGs, using a consensus bootstrap approach (Methods). Of the 465 DEGs, 405 were assigned to 13 named modules (Fig 4A), of which five had stable bootstrap support (LFP *>* 0.70; Supplementary Fig S8). Gene Ontology enrichment confirmed the modules as biologically coherent (Supplementary Fig S9). The pink module, which contains *jmj2* and *jmj4*, was enriched for ribosome biogenesis, negative regulation of gene expression (epigenetic), and positive regulation of growth, with the epigenetic and growth terms driven by the JMJ cluster itself (Supplementary Table S11). The greenyellow module was enriched for developmental cell growth and DNA replication, and contains *sec6* (log_2_ FC = +2.87), an exocyst component identified as a plant height GWAS candidate (Peiffer *et al*. 2014), and *pcna2* (log_2_ FC = +2.99), a DNA-replication factor (Supplementary Table S10). The purple module contained the plant height GWAS candidate *exo*/*phi1* (log_2_ FC = *−*2.39; Table 1) and was enriched for brassinosteroid signaling regulation and negative regulation of chromatin silencing.

**Table 1.**
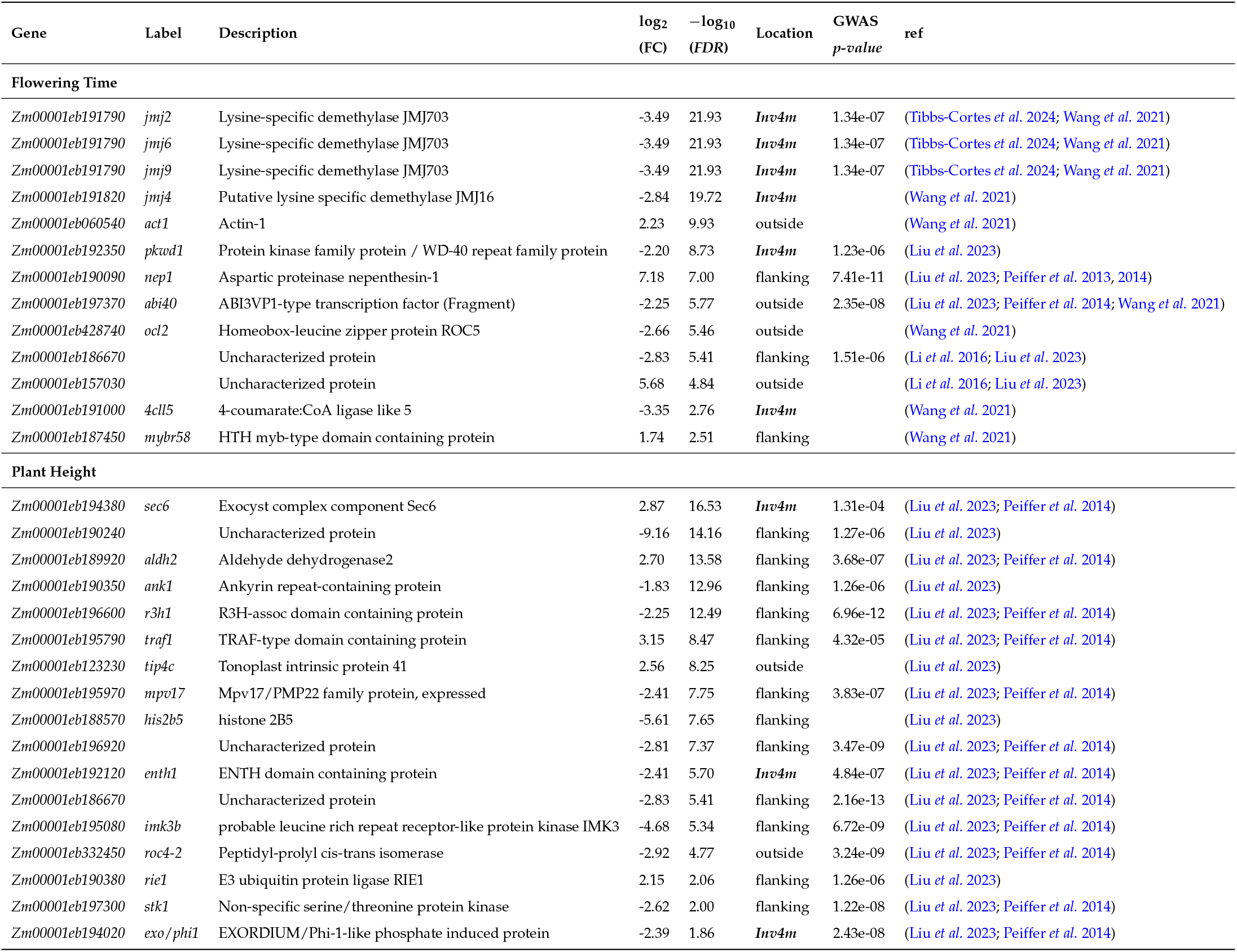
Effect of *Inv4m* on Flowering Time and Plant Height Gene Candidates.

**Figure 4.**
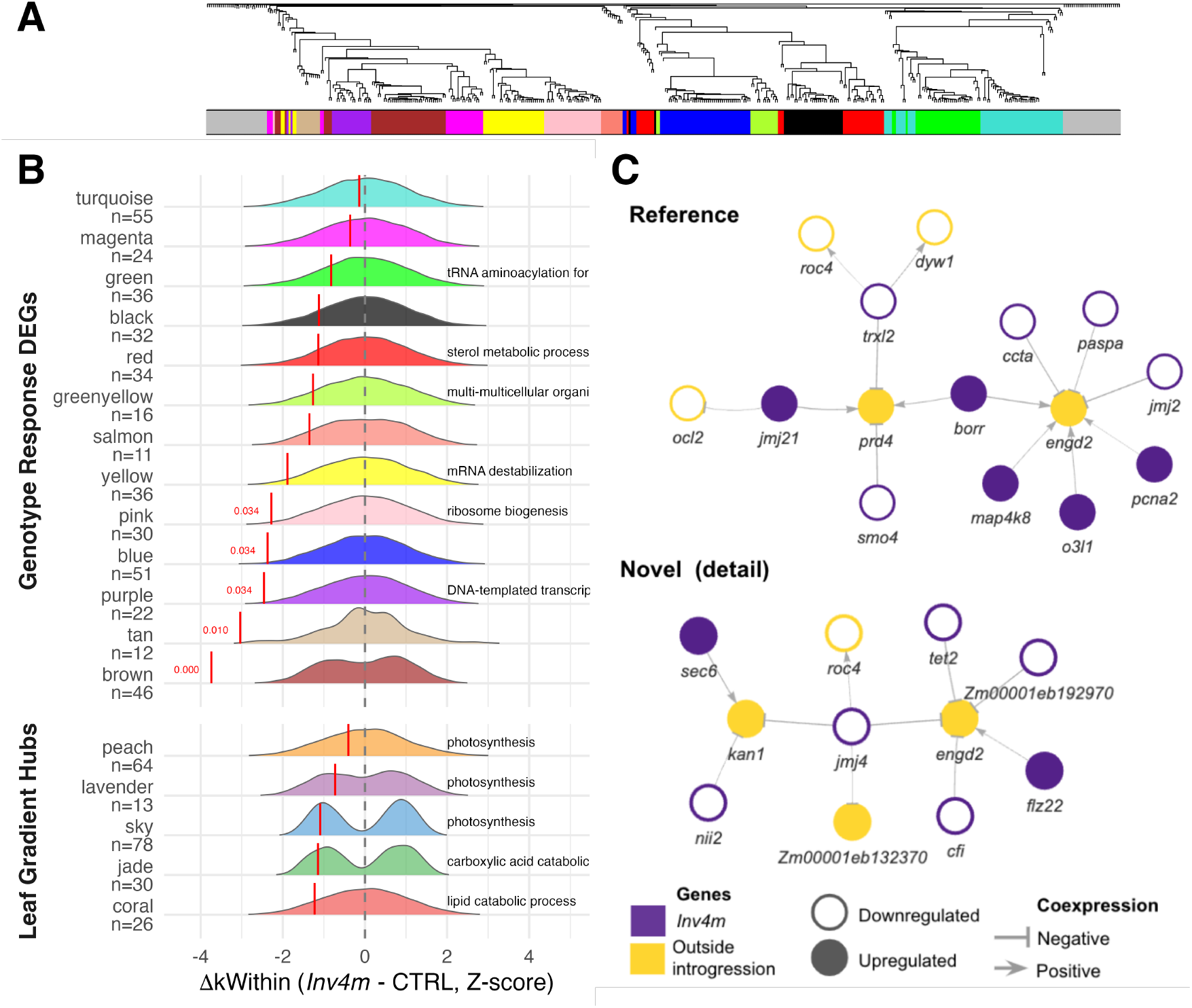
Network analysis of *Inv4m* DEGs. **(A)** Consensus WGCNA modules of *Inv4m* DEGs reconstructed from CTRL plant samples only. The 13 gene coexpression modules were obtained from 1000 consensus iterations with 80% gene subsampling and variable *minModuleSize* (10 to 35). The color bar indicates module membership for 465 DEGs (FDR *<* 0.05). **(B)** Change in intramodular connectivity (Δ*k*_*Within*_) of the genes in each module from (A) between *Inv4m* and CTRL plants. Density ridges show the null distribution of *z*-scored Δ*k*_*Within*_ obtained by shuffling genotype labels across 1000 permutations; red vertical segments mark the observed *z*-score for each module. Modules are grouped by gene set: Genotype Response DEGs (top facet) and Leaf Gradient Hubs (bottom facet). FDR adjusted *p*-values are shown in red next to modules with significant connectivity loss. Text to the right of each ridge shows the top GO Biological Process term for each module. **(C)** Trans coexpression network from *Inv4m* strong DEGs, reconstructed jointly from CTRL and *Inv4m* samples and shown as minimum spanning trees. *Reference* edges recover coexpression relationships reported in MaizeNetome. *Novel* edges (detail) show the *jmj4* neighborhood subgraph; the full Novel network is in Supplementary Fig S11. Nodes within the *Inv4m* introgression are purple and targets elsewhere in the genome are gold. Node fill indicates direction of differential expression: filled nodes are upregulated and open nodes are downregulated in *Inv4m* plants. Edge polarity reflects the sign of mutual information: arrows indicate positive coexpression and bars indicate negative coexpression.

We then applied a genotype label permutation test on intramodular connectivity (Δ*k*_*Within*_; Methods) to test whether modules show genotype specific connectivity loss beyond sample to sample variability. Five of 13 modules reached significance (brown, tan, purple, blue, pink; FDR *<* 0.05; Fig 4B; Supplementary Table S8), with brown showing the largest effect. As a topological control, we repeated the entire pipeline on an orthogonal “Leaf Gradient” reference network built from genes varying along the leaf developmental gradient. None of its five modules reached significance under the same permutation test (Fig 4B, bottom facet; Supplementary Table S9), isolating the connectivity loss as a *Inv4m* specific effect rather than a genome wide attenuation of coexpression. The pink module illustrates the pattern. *jmj2* is among the pink module’s top hubs; both *jmj2* and *jmj4* had high *k*_*Within*_ in CTRL (*∼*3.3 and *∼*3.2 respectively) and dropped to*∼* 0.6 in *Inv4m* (Supplementary Fig S10). The exocyst component *sec6*, by contrast, went from *k*_*Within*_ = 0.81 in CTRL to effectively zero in *Inv4m*, suggesting that its upregulation decouples it from native coexpression partners.

### Trans regulatory network analysis detects potential Inv4m epigenetic regulators belonging to a growth related module

To investigate the potential trans regulatory effects of the *Inv4m* chromosomal inversion, we constructed a coexpression network linking DEGs within the introgressed region (potential regulators) to differentially expressed genes located elsewhere in the genome (trans targets). We validated these coexpression relationships against the MaizeNetome database, which catalogs experimentally-supported regulatory interactions in maize (Fig 4C).

The trans regulatory network comprised 135 genes connected by 552 coexpression edges. Of the 103 potential regulator genes within or flanking the introgression, 33 were located within the *Inv4m* proper and 70 in the flanking regions. These potential regulators showed coexpression relationships with 32 trans target genes distributed across all ten maize chromosomes. We classified 16 edges (2.9%) as “Reference” because they appear in the MaizeNetome database, and the remaining 536 edges (97.1%) as “Dataset-specific.” The low overlap likely reflects differences in tissue representation and developmental context: MaizeNetome aggregates expression from 7,603 RNA-seq samples spanning diverse tissues and genotypes (Feng *et al*. 2023), whereas our network captures coexpression relationships specific to field grown V10-V12 leaves. The dataset specific edges are statistically well supported in our experiment (Pearson correlation *FDR <* 0.05) and their absence from MaizeNetome indicates condition specificity rather than lack of biological support.

The Reference network included known regulatory relationships involving *proliferating cell nuclear antigen2* (*pcna2*; log_2_ *FC* = +2.99), a DNA replication factor, and *outer cell layer2* (*ocl2, Zm00001eb428740*; log_2_ *FC* = *−*2.66), a HD-ZIP IV transcription factor involved in epidermal cell differentiation. Notably, the Reference network also contained edges connecting *jmj2* (log_2_ *FC* = *−*3.49) to trans targets, supporting a conserved regulatory role for this histone demethylase.

The dataset specific network identified a previously uncharacterized *jmj4* neighborhood subgraph (Fig 4C). *jmj4* (log_2_ *FC* = *−*2.84) showed strong coexpression with *Exocyst complex component SEC6* (*sec6*; log_2_*FC* = +2.87), which functions in vesicle trafficking and cell plate formation during cytokinesis. This connection suggests that the JMJ cluster may influence cell division through pathways beyond direct chromatin modification of target gene promoters. The enrichment of growth regulation terms among *Inv4m* downregulated genes is consistent with the altered SAM geometry observed in *Inv4m* plants, suggesting that perturbation of epigenetic regulators may underlie the developmental phenotypes.

### JMJ cluster expression differences mainly reflect copy number variation

The *jmj2*/*jmj4* cluster exhibited the strongest transcriptional response to *Inv4m* introgression, with *jmj2* showing log_2_ FC = *−*3.49 (*FDR <* 10^*−*20^), the most significant of all DEGs. This apparent downregulation was consistent across all 13 tissue experiment combinations examined, including both PA2022 field samples (V13 adult leaves) and Crow et al. (2020) growth chamber samples spanning roots, leaves, and shoot apical meristems (Fig 5A).

**Figure 5.**
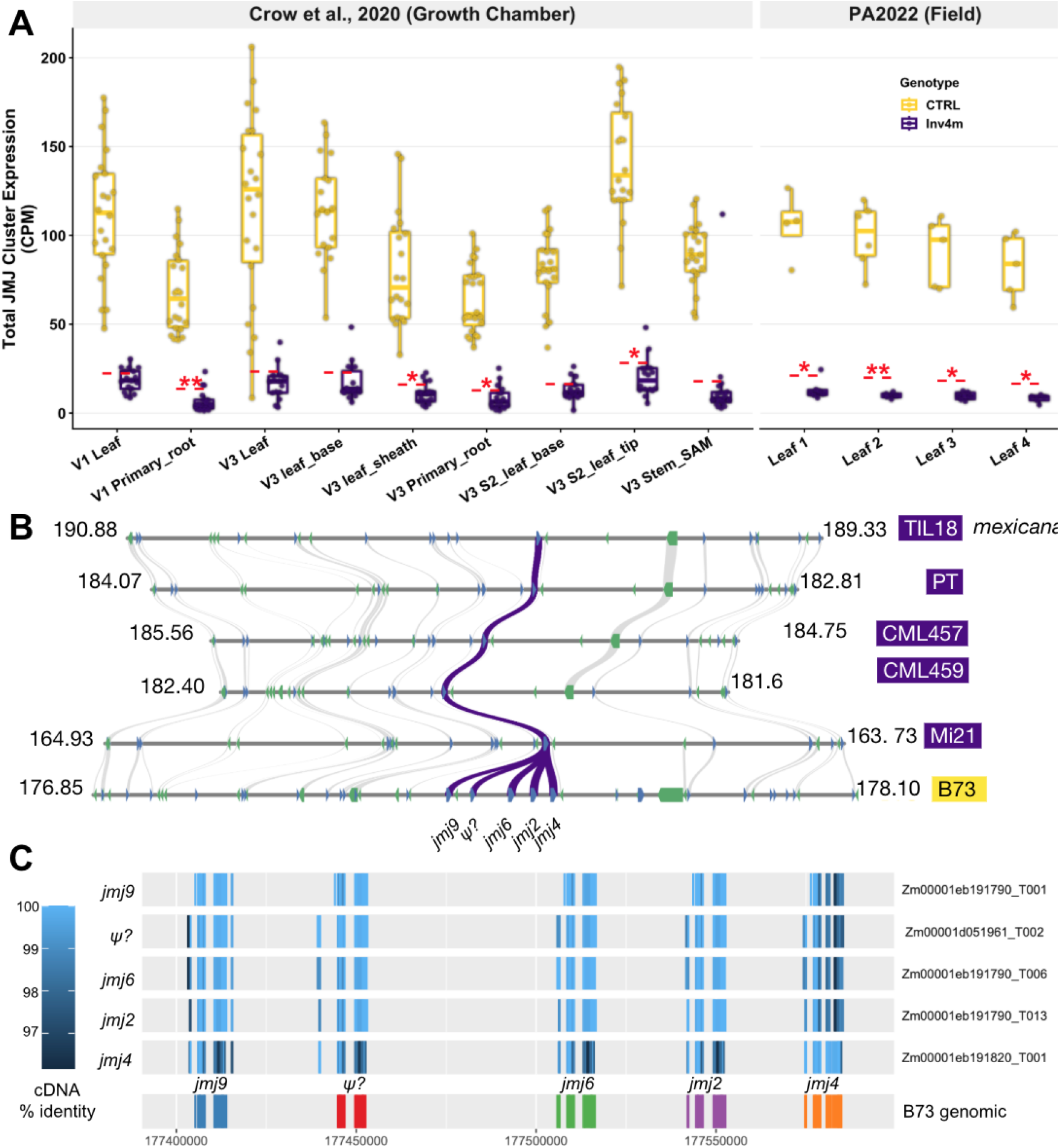
The *jmj2*/*jmj4* cluster is constitutively downregulated in *Inv4m* and shows lineage specific expansion in B73. **(A)** Total JMJ cluster expression (sum of *jmj2* + *jmj4*) across tissues and experiments. Boxplots show CPM expression in CTRL (gold) and *Inv4m* (purple) genotypes. Left: Crow et al. (2020) growth chamber data across 9 tissue types. Right: PA2022 field data across 4 leaf positions. Dashed red line indicates expected *Inv4m* expression under copy number scaling (CTRL/5), assuming the single remaining *jmj4* copy produces 1/5 of the total cluster output. Red asterisks denote significant deviation from expectation (linear contrast *t*-test, FDR *<* 0.05: *, *<* 0.01: **, *<* 0.001: ***). **(B)** Microsynteny of the JUMONJI gene cluster across five *Zea* genomes. From top to bottom: the teosinte TIL18 (*Z. mays* ssp. *mexicana*), the highland landrace Palomero Toluqueño (PT), and the CIMMYT highland inbreds CML457 and CML459 each contain a single *jmj4* ortholog, while B73 (bottom) has an expanded tandem array of five paralogs (*jmj9, ψ*?, *jmj6, jmj2, jmj4*). Purple ribbons indicate syntenic relationships between *jmj* loci; gray ribbons show conserved flanking genes. Coordinates in Mb. **(C)** Transcript structure and annotation of the B73 *jmj* cluster. The NAM v5 annotation consolidates *jmj2, jmj6*, and *jmj9* as alternative transcripts of a single locus (*Zm00001eb191790*), while *jmj4* is annotated separately (*Zm00001eb191820*). The putative pseudogene (*ψ*?) retains only a v4 identifier (*Zm00001d051961*). Heatmap intensity indicates cDNA sequence identity (97 to 100%). Bottom track shows genomic positions of individual cluster members.

However, microsynteny analysis across five *Zea* genomes showed that the B73 reference contains a tandem expansion of five JMJ paralogs (*jmj9*, a putative pseudogene, *jmj6, jmj2*, and *jmj4*), whereas highland teosinte TIL18, the highland landrace Palomero Toluqueño, and the CIMMYT tropical inbred lines CML457 and CML459 each retain only a single *jmj4* ortholog (Fig 5B). The single copy state is shared across highland and tropical backgrounds, suggesting it represents the ancestral configuration. The high sequence identity among B73 paralogs (*>*97% cDNA similarity; Fig 5C) suggests recent duplication and complicates individual paralog quantification.

To test whether the observed expression differences could be explained by copy number alone, we compared total cluster expression in *Inv4m* plants against the expectation under simple dosage scaling (CTRL/5). In field samples, *Inv4m* expression was consistently and significantly below this expectation: Leaf 1 achieved 61.5% of expected levels (*t* = 2.94, *FDR* = 0.031), while Leaves 2 to 4 showed only 49 to 53% of expected expression (*FDR <* 0.015 for all). This pattern suggests that beyond copy number reduction, additional regulatory mechanisms suppress JMJ cluster transcription in the field.

In contrast, growth chamber data showed a less consistent pattern. Only 4 of 9 tissues exhibited significant deviation from copy number expectation after FDR correction, with expression levels ranging from 48% (V1 root) to 83% (V1 leaf) of expected values. Notably, the shoot apical meristem showed no significant deviation (81.8% of expected, *FDR* = 0.57). This environment dependent pattern may reflect condition specific regulatory mechanisms acting on the JMJ cluster beyond the baseline copy number effect.

## Discussion

Megabase-scale inversions are repeatedly implicated in local adaptation across taxa, but the specific genes through which they drive their phenotypic effects have remained largely unidentified (Wellenreuther and Bernatchez 2018b; Kirkpatrick and Barton 2006; Kirkpatrick 2010). Here, we used near isogenic lines, multi-environment field experiments, comparative genomics across five *Zea* genomes, and gene network analysis to dissect *Inv4m*, a 15 Mb inversion introgressed into maize from *mexicana*. We found that, compared with the B73 temperate reference, which carries a cluster tandem repeat of five JUMONJI histone demethylase paralogs, the highland materials carrying *Inv4m* only retain a single copy. This single copy of the JMJ demethylase is part of a gene regulatory network that may explain *Inv4m*’s effects on flowering time acceleration and growth and its adaptive value. Overall, we identify a candidate adaptive locus within *Inv4m*, suggest a regulatory mechanism by which a single dosage sensitive hub can shape an inversion encoded phenotype, and provide a worked example of how single reference genome inference can mistake the derived for the ancestral state at adaptive loci.

### Repeat-mediated breakpoints suggest an ectopic recombination origin

The breakpoint architecture we observe at *Inv4m*, asymmetric *knob180* and *TR-1* tandem arrays whose size, content, and orientation are mirrored between standard and inverted karyotypes, has the hallmarks of an ectopic recombination origin. Under this mechanism, mispairing between dispersed repeat copies during meiosis generates rearrangements whose breakpoints reside within the participating repeats (Zhang and Peterson 2004; Delprat *et al*. 2009), and the resulting inversion swaps the two breakpoint segments, producing exactly the mirror image pattern we see between karyotypes.

This pattern is not unique to *Inv4m*. The maize abnormal chromosome Ab10 carries two megabase scale inversions, each preceded by *TR-1* repeat arrays (Liu *et al*. 2020; Mroczek *et al*. 2006), and the inbred line Zi330 displays inverted knob band orders on homologous chromosomes, implying recurrent knob-mediated rearrangement (Yang and Xu 2012). In *Arabidopsis thaliana* Col-0, the 750 kb *hks4* heterochromatic knob overlaps a 1.2 Mb paracentric inversion relative to Ler and *A. lyrata* (Schmidt *et al*. 2020; Fransz *et al*. 2000; Zapata *et al*. 2016). The pattern extends well beyond plants: in deer mice (*Peromyscus maniculatus*), 21 megabase scale inversion polymorphisms are flanked by long inverted repeats and segmental duplications mapping to gene-poor centromeric satellite arrays (Harringmeyer and Hoekstra 2022; Gozashti *et al*. 2025). Further, in the Atlantic cod (*Gadus morhua*) megabase scale inversions display two asymmetric breakpoint segments, the closest structural parallel to *Inv4m* we have identified (Aasegg Araya *et al*. 2025). Across these systems, repeat-rich gene-poor regions appear to serve both as the substrate for inversion generating ectopic recombination and as locations where such rearrangements can occur without disrupting genes.

Two questions our data cannot resolve are when the *Inv4m* breakpoints formed and which specific repeat copies participated. The repeat arrays themselves carry no clear molecular clock, and recombination suppression within the inversion preserves the founding sequence, blocking direct inference of timing. Whether the cross taxon parallels imply a shared mechanism at the root of *Inv4m* origin therefore remains a working hypothesis rather than a demonstrated history.

### Genomic and functional evidence suggest a link between Inv4m jmj2/jmj4 copy number, growth, and flowering time plasticity

Among the 465 genes differentially expressed between *Inv4m* and CTRL plants, a cluster of JUMONJI histone demethylases provides the most consistent signal across our field data and the independent growth chamber experiments of Crow *et al*. (2020) (Fig 5A). Microsynteny across five *Zea* genomes show that B73 carries five paralogs, while highland teosinte (TIL18), the highland landrace Palomero Toluqueño, and the CIMMYT highland inbreds CML457 and CML459 each retain a single *jmj4* ortholog (Fig 5B). The 5:1 copy number difference accounts for most of the apparent expression fold change (log_2_ FC = *−*3.49; Table 1).

Plant histone methylation affects gene expression and operates through opposing systems finely regulated by methylation writing and erasing enzymes. Generally, histone methylation is associated with downregulated gene expression. However, H3K4 trimethylation is known to increase gene expression. This methylation mark is removed by members of the KDM5/JARID1 subfamily, to which the *Inv4m jmj2*/*jmj4* demethylase cluster belongs (Qian *et al*. 2019). On the other hand, H3K9 dimethylation functions as a repressor and is removed by demethylases of the JHDM2/KDM3 subfamily Hung *et al*. (2021); **?**, to which *jmj21* belongs. In *Inv4m, jmj2*/*jmj4* is downregulated due to copy number variation, and *jmj21* is upregulated. Therefore, their combined effects should lead to higher expression of their target genes.

The direction of these phenotypic effects is harder to predict because it depends on whether the affected target genes act as activators or repressors. Functional studies across grasses and Arabidopsis link the KDM5/JARID1 subfamily to both flowering time and growth, but in a target- and context-dependent direction. In rice, *OsJMJ703* knockdown accelerates flowering (Song *et al*. 2018). In Arabidopsis, JMJ14 demethylates H3K4me3 at *FLOWERING LOCUS T*, so JMJ14 loss of function mutants flower early (Lu *et al*. 2010; Yang *et al*. 2012b). Yet overexpression of the related JMJ15 also accelerates flowering by demethylating the floral repressor *FLC* (Yang *et al*. 2012a).

The JHDM2/KDM3 H3K9 demethylase family, which includes *jmj21*, has been shown to regulate flowering: overexpression of Arabidopsis JMJ28 accelerates flowering by removing H3K9me2 at *CONSTANS* to activate *CO* (Hung *et al*. 2021). While our study suggests that similar mechanisms may underlie flowering time acceleration due to *Inv4m* JMJ demethylases, further confirmation of the specific target genes would support this hypothesis. However, independent GWAS evidence in maize links the JMJ cluster region directly to phenotypic plasticity of flowering time: two SNPs near *jmj9* are associated with the slope of days to silking across environments (Tibbs-Cortes *et al*. 2024).

Differential regulation of methylation patterns of H3K4 and H3K27 also affects the expression patterns of cytokinin metabolism (Chen *et al*. 2013; Bellegarde *et al*. 2025) and regulates internode length and plant height. However, we do not observe significant changes of genes involved in cytokinin synthesis or degradation.

Due to the epigenetic regulatory nature of JMJ genes, its effect is interlinked with changes we observed in our coexpression network analysis. Of thirteen coexpression modules built from the 465 DEGs, five show genotype-specific connectivity loss in *Inv4m* under a label permutation test (Fig 4), including the pink module which includes the JMJ cluster. The pink module is one of the five with significant module-level connectivity loss in the permutation test; both genes had *k*_*Within*_ *∼* 3.2 in CTRL and *∼* 0.6 in *Inv4m*.

A *trans*-coexpression network independently connects the JMJ cluster to growth related targets including *pcna2*, a DNA replication factor, and *sec6*, an exocyst component and plant height GWAS candidate (Peiffer *et al*. 2014), both upregulated in *Inv4m* (Fig 4C). Two additional WGCNA modules reinforce the growth network interpretation: the greenyellow module includes *sec6* and *pcna2* together with the cell polarity regulator *ropgef14*; the purple module includes *exo*/*phi1* (EXORDIUM/Phi-1-like), another plant height GWAS candidate (Liu *et al*. 2023; Peiffer *et al*. 2014) that mediates brassinosteroid responsive cell expansion in Arabidopsis (Farrar *et al*. 2003; Schröder *et al*. 2009).

### Limitations of the study

The introgression retains 24 Mb of flanking sequence around the 15 Mb inversion (reduced from 57 Mb in earlier BC_5_ populations (Crow *et al*. 2020)), so attribution to *Inv4m* itself rather than the broader haplotype cannot be made from this experiment alone. The *trans*-coexpression analysis partly addresses this by restricting candidate regulators to within the 15 Mb inversion, and the ZeaL panel replicates the flowering effect across multiple donor backgrounds; nonetheless, definitive fine mapping will require restoring recombination within *Inv4m*, an approach now made feasible by CRISPR-engineered inversion reversal in maize (Schwartz *et al*. 2020).

The high sequence identity among B73 JMJ paralogs (>97%) complicates individual paralog quantification. To establish whether the JMJ dosage reduction drives the network reorganization we observe, or whether both reflect parallel consequences of the same structural variant, direct tests would need to be performed. These include ChIP-seq for H3K4me3 and H3K9, the targets of the differentially expressed JMJ demethylases, across *Inv4m* karyotypes, transgenic manipulation of JMJ copy number, and Hi-C analysis of chromatin organization on chromosome 4. The B73-specific expansion of JMJ paralogs also carries a methodological lesson beyond *Inv4m*. When an inversion is interpreted by aligning haplotypes to a single reference, the polarity of structural variation can be misread: apparent downregulation of genes within the inversion may simply reflect copy number variation. This caveat applies broadly to eQTL studies and to inversion analyses across taxa, and it is straightforwardly addressed by including multiple reference-quality assemblies spanning the standard and inverted haplotypes, as in the comparative genomics design we used here.

A related but distinct caveat applies to our differential expression analysis. Sequence divergence between B73 and Mi21 inside *Inv4m* (median CDS divergence *∼*1.1%) raises the possibility that some apparent expression differences at the most diverged genes reflect reduced read mapping for diverged transcripts rather than regulatory change.

### Conclusions

Two classical theories address how inversions become adaptive: the supergene hypothesis (Dobzhansky 1971), invoking epistasis among captured alleles, and the local adaptation hypothesis (Kirkpatrick and Barton 2006; Kirkpatrick 2010), invoking recombination suppression of locally favored allele combinations against gene flow. *Inv4m*’s elevation-dependent fitness reversal in Mexican landraces (Crow *et al*. 2020) fits the local adaptation framework; the supergene hypothesis is difficult to test directly because recombination suppression precludes the experimental dissection of epistasis. Our results suggest a mechanism that both theories implicitly assume but neither explicitly states: some of *Inv4m*’s phenotypic effects may flow through a single dosage-sensitive regulator whose action propagates through a downstream gene network extending beyond the inversion. This perspective is consistent with cross-taxon evidence that adaptive inversions reshape gene expression beyond what sequence divergence alone predicts.

Recombination suppression would then protect a co-adapted regulatory architecture, not just a collection of independent alleles.

## Materials and Methods

### Inv4m near isogenic lines, growth conditions, experimental design, and phenotype measurements

To measure the effects of the *Inv4m* in plant field phenotypes and their phosphorus starvation response transcriptome, we used a highland traditional variety carrying the Highland haplotype of *Inv4m* corresponding to the inverted karyotype. The accession Michoacán 21 (referred to as Mi21), from the Mexican Cónico group, was obtained from the International Maize and Wheat Improvement Center (CIMMYT). In contrast, the reference genome of the temperate inbred B73, the recurrent parent for introgression, carries the lowland haplotype corresponding to the standard non-inverted karyotype at *Inv4m*.

From the cross of Mi21 with B73 one F1 individual was back-crossed to B73 for eight generations. We selected lines carrying *Inv4m* with a diagnostic SNP during each cycle using a cleaved amplified polymorphic sequence (CAPS) marker. The marker SNP is PZE04175660223 located at position 4:181637780 in the NAM B73v5 *Zea mays* genome assembly. Amplification of the polymorphic site was done with the following primer pair: *CT-GAGCAGGAGATGATGGCCACTC* and *GGAAAGGACATAAAA-GAAAGGTGCA*, and subsequently cleaved by *HinfI*. Plants were genotyped using the CAPS marker for selecting heterozygous plants at BC8S2 after selfing seeds of *Inv4m* and CTRL homozygous individuals were selected for the field trial.

Plants were planted on May 26, 2022 at the Russell E. Larson Agricultural Research Farm in Rock Springs, Pennsylvania (40°42’36” N 77°57’0” W, 366 masl) in soil classified as a Hagerstown silt loam (fine, mixed, semiactive, mesic Typic Hapludalf). Experimental conditions were similar to those previously described in (Strock *et al*. 2018). The experiment had a complete block design with two phosphorus (P) levels. Low-P fields (5 ppm Mehlich-3 Phosphorus) and high-P fields (36 ppm Mehlich-3 Phosphorus) were divided into smaller blocks. Three rows per block were planted with a mean stand count of 8 plants per plot, and the plants from the center row were selected for measurements to avoid border effects. Fields received fertilization based on treatment requirements. Drip irrigation was provided during dry periods. Each genotype was replicated four times within its P treatment.

### Phenotype analysis

For stover mass growth curves, a different plant at each time point 40, 50, 60, and harvest, 121 days after planting (DAP), was collected, dried, and weighed for the same row. Stover dry mass data was fitted to a logistic growth model using the R package *Growthcurver* (Sprouffske and Wagner 2016). Maximum stover dry weight was estimated as the maximum over the four time points rather than dry weight at harvest. Ear measurements were taken for one ear per row at harvest.

We used a two stage approach to estimate *Inv4m* effects on field phenotypes while accounting for spatial heterogeneity. In the first stage, we applied P-spline analysis of spatial trends (SpATS) (Rodríguez-Álvarez *et al*. 2018) implemented via the `statgenHTP::fitModels()` function (Millet *et al*. 2025). For each phenotype *y*, SpATS fits a mixed model that decomposes spatial variation into smooth bivariate surfaces using two dimensional P-splines over field row and column coordinates:

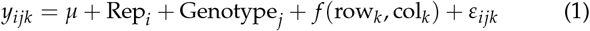

where *µ* is the overall mean, Rep_*i*_ is the replicate effect, Genotype_*j*_ is the genotype effect, and *f* (row_*k*_, col_*k*_ ) is a smooth bivariate spatial surface estimated via penalized spline ANOVA (PSANOVA). Phosphorus treatment is *not* included in this stage so its variance remains in the corrected values for proper modeling in the second stage. For the PA2024 hybrid trial, lines were treated as a random effect cohort and the two hybrid entries Inv4_`{B73, Mi21}` as fixed effects via addCheck = TRUE. Per-plot spatially corrected values were extracted with `getCorrected()` from the resulting fit.

In the second stage, we tested for *Inv4m* effects on the spatially corrected values using a linear model with genotype and phosphorus treatment as fixed factors:

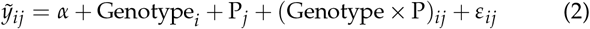

where 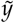 denotes a spatially corrected per-plot value. Marginal Genotype contrasts (*Inv4m −*CTRL, averaged over phosphorus levels) and conditional Genotype contrasts within each phosphorus level were estimated using the emmeans package (Lenth and Piaskowski 2025). For each trait this yields three contrasts (one marginal, two conditional), all of which are pooled into a single Benjamini-Hochberg false discovery rate family across the (trait*×* contrast) table (Benjamini and Hochberg 1995). Significance stars in Figure 2 panels A and B reflect the marginal-Genotype row. The Genotype*×* phosphorus interaction *p*-value is retained as a model diagnostic and is not included in the FDR pool.

Days to anthesis (DTA) and days to silking (DTS) were converted to growing degree days (GDDA and GDDS, respectively) using daily temperature records from the Rock Springs weather station with a base temperature of 10°C and an upper threshold of 30°C.

### Genotype × Environment analysis

To assess the consistency of *Inv4m* effects across environments, we analyzed phenotype data from three location-year combinations: Pennsylvania 2022 (PA2022), Pennsylvania 2025 (PA2025), and North Carolina 2025 (NC2025). All environments used the Mi21 donor background. Phenotypes analyzed included plant height (PH), days to anthesis (DTA), days to silking (DTS), and their thermal time equivalents (GDDA, GDDS) calculated using growing degree days with a base temperature of 10°C and an upper threshold of 30°C.

Spatially corrected phenotype values were obtained for each environment using SpATS as described above. We then performed two way ANOVA with genotype (*Inv4m* vs. control) and environment as fixed factors:

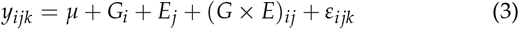

where *G*_*i*_ is the genotype effect, *E*_*j*_ is the environment effect, and (*G × E*)_*ij*_ is the genotype by environment interaction. Effect sizes were quantified using partial eta squared 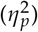 for ANOVA effects and Cohen’s *d* for pairwise genotype comparisons within each environment. Marginal means and contrasts were estimated using the *emmeans* package (Lenth and Piaskowski 2025). P-values were adjusted for multiple testing using the Benjamini-Hochberg FDR method (Benjamini and Hochberg 1995).

To examine temperature sensitivity of *Inv4m* effects, we correlated genotype effect sizes (Cohen’s *d*) with mean environmental temperature during the first 100 days after planting, calculated from NASA POWER hourly temperature data.

### ZeaL NIL panel Inv4m effect (NC2025)

To test whether the *Inv4m* flowering time effect generalizes beyond the Mi21 single donor background, we analyzed the Zea Exotic Allele Library (ZeaL) NIL panel grown in 2025 at the Central Crops Research Station (Clayton, NC). The ZeaL panel introgresses chromosomal segments from multiple teosinte donor accessions into the B73 background. We restricted the analysis to BC1 families segregating at the *Inv4m* locus, that is, families containing at least one plant carrying the donor (teosinte) *Inv4m* inverted karyotype and one plant carrying the recurrent (B73) standard karyotype, so that the *Inv4m* fixed effect could be estimated from a within family contrast. This filter retained four *Z. mays* ssp. *mexicana* ancestry groups (Chalco, Durango, Mesa-Central, Nobogame) and *Z. mays* ssp. *huehuetenanguensis* (Huehuetenango), and excluded donor lineages that do not carry the *Inv4m* inverted karyotype (Balsas parviglumis, *Z. luxurians, Z. diploperennis*) because every plant from those lineages was monomorphic standard at *Inv4m*. Our ancestry groups correspond to the teosinte races defined by (Sánchez González *et al*. 2018); the column is stored as `race` in the ZeaL germplasm meta-data and renamed to `ancestry` on join.

Spatially corrected phenotypes (DTA, DTS, PH) were obtained as described above. Three known outlier plots (IDs 2756, 453, 304) were removed; the remaining dataset comprised 393 plots from 218 NILs across 35 BC1 families. Pedigree levels (donor accession, F1, BC1, BC2) were derived as substring prefixes of the NIL identifier; ancestry was joined from the ZeaL germplasm metadata via donor accession. For each trait we fit a linear mixed model with *Inv4m* as a two level fixed factor (recurrent vs. donor) and a strictly nested random effect structure spanning ancestry, BC2 family, and NIL identity:

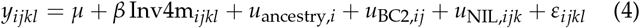

Models were fit with `lme4` (Bates *et al*. 2014) via `lmerTest` (Kuznetsova *et al*. 2017) for Satterthwaite degrees of freedom. Intermediate pedigree levels (donor accession, F1, BC1) carry zero additional variance once ancestry (above) and BC2 (below) are accounted for, so we report the parsimonious 3 random effect specification; the full slash nested 6 random effect model gave identical *Inv4m* fixed effect estimates with singular fits at the intermediate levels. We additionally validated the parametric inference with a within BC2 permutation test (1000 permutations of *Inv4m* labels within each BC2 family); permutation *p* values agreed with Satterthwaite *p* values within rounding. Per trait *p* values were adjusted for multiple testing across the 3 trait family (DTA, DTS, PH) using the Benjamini Hochberg method (Benjamini and Hochberg 1995). For visualization (Supplementary Fig S2), lineage corrected trait values were computed by subtracting the ancestry and BC2 random effect BLUPs but retaining the NIL random effect, 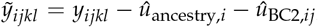 so that the within arm spread reflects the line to line NIL variation the model uses for inference. The reproducible analysis notebook is at `scripts/inversion_paper/Zeal_Inv4m_flowering_lmm.Rmd`.

### Internode length measurements

To characterize the effect of *Inv4m* on plant architecture, we measured internode lengths in the NC2025 field experiment. At physiological maturity, plants were harvested and internodes were measured sequentially from the tassel bearing node (position 1) downward. Internode position was recorded both as absolute node number (from ground level) and as position from the top (from tassel). The ear bearing node was marked for each plant.

Internode profiles were analyzed separately for each donor background (Mi21 and TMEX). For the TMEX background, measurements were limited to the top 10 internodes due to plant architecture differences. Within each donor, we compared *Inv4m* and control genotypes using two sample *t*-tests for total node count. Internode length profiles were visualized by plotting length against position from top, with individual plant trajectories shown.

To validate the internode measurements against whole plant phenotypes, we compared the sum of measured internode lengths with directly measured plant height (PH) from field records using linear regression. Agreement between methods was assessed by *R*^2^ and slope deviation from unity.

### Mi21 NIL genome assembly and scaffolding

To obtain a reference-quality genome for the *Inv4m* donor parent haplotype, we performed *de novo* assembly of the Mi21 near isogenic line from PacBio HiFi sequencing data. Two barcoded HiFi libraries (bc2051, *∼* 9*×* coverage; bc2052, *∼* 5*×* coverage) were combined for a total of *∼* 14*×* genome coverage. Primary contigs were assembled with hifiasm v0.25.0 (Cheng *et al*. 2021b) using the -l0 flag to disable purge duplication, yielding 740 contigs totaling 2.21 Gb with a contig N50 of 13.7 Mb.

Contigs were scaffolded against two reference genomes using RagTag (Alonge *et al*. 2022): Zm-B73-REFERENCE-NAM-5.0 (Hufford *et al*. 2021) as the recurrent parent background and Zm-PT-REFERENCE-HiLo-1.0 as a highland representative carrying the standard *Inv4m* arrangement. Each scaffolding was performed with `ragtag.py scaffold –u` (unplaced contigs retained) using minimap2 (Li 2018) alignments. The two resulting AGP files were merged with `ragtag.py merge`, producing 10 pseudochromosomes and 455 unplaced contigs. The merged scaffold placed 97.6–98.5% of assembled bases onto chromosomes.

### Gene model transfer

Gene models were transferred from B73 (Zm00001eb.1 annotation) to the Mi21 NIL assembly using Liftoff (Shumate and Salzberg 2021) with parameters `-s 0.5 -a 0.5 -flank 0.1 -cds –copies`. This lifted 1,145,935 features, with 252 features unmapped. The `-copies flag` was used to detect copy number variation at the JUMONJI histone demethylase cluster on chromosome 4.

### Chromosomal knob repeat annotation

To characterize the repeat content of the *Inv4m* breakpoint segments across genomes, we performed BLAST+ (Camacho *et al*. 2009) similarity searches of chromosome 4 in B73 (NAM5), PT (HiLo-1.0), TIL18 (PanAnd-1.0), and the Mi21 NIL assembly using representative sequences of two classes of chromosomal knob repeats as queries. For the 180 bp knob repeat we used accession M32524.1 (Dennis and Peacock 1984), and for the *TR-1* repeat we used accession AY083942.1 (Hsu *et al*. 2003). Searches were run with the `megablast task (blastn -task megablast)`, which uses a word size of 28 and is optimized for highly similar sequences. Hits were plotted along chromosome 4 with normalized bitscore on the *y*-axis (Fig 1B). Breakpoint annotation coordinates were defined by the AnchorWave (Song *et al*. 2022) genome alignment endpoints where reference-orientation synteny was lost (Supplementary Table S2).

For breakpoint self-similarity analysis, we extracted the unaligned segments flanking each breakpoint and performed local self-alignment with LAST (Kiełbasa *et al*. 2011). The resulting dotplots (Fig 1D) show tandem repeat arrays as diagonal and off diagonal patterns, with forward matches shown in red and reverse (inverted) matches in blue.

### Whole-genome alignment and microsynteny

To confirm the orientation of *Inv4m* in the Mi21 NIL assembly and delimit breakpoint coordinates, we performed CDS-anchored whole genome alignments against B73, PT, and TIL18 using AnchorWave (Song *et al*. 2022) `proali with -R 1 -Q 1` (one to one mapping, no whole genome duplication). CDS anchor sequences were extracted from each reference GFF3 with `anchorwave gff2seq` and mapped to both reference and query assemblies using minimap2 (Li 2018) (`-x splice -k 12 -p 0.4 -N 20`). CDS-anchor dotplots were filtered to uniquely mapped anchors (MAPQ *≥*60) and dominant scaffold pairings to reduce multi mapping noise from duplicated gene families (Fig 1C).

To examine microsynteny at the JMJ cluster, we performed pairwise ortholog detection between B73 and the Mi21 NIL assembly using jcvi/MCScan (Tang *et al*. 2008) with LAST (Kiełbasa *et al*. 2011) alignments `(–cscore=.99, –minspan=30)`. The resulting synteny blocks confirmed that all B73 JMJ paralogs at the locus map to a single Mi21 gene (Mi21_Zm00001eb191820), consistent with a single copy state in the inverted haplotype (Fig 5B).

### Tissue sampling, RNA extraction, and sequencing

We sampled the plants at 63 DAP when we estimated them to be between v10 to v12 developmental stages. We took tissue from the first leaf with a fully developed collar, or first leaf before the flag leaf, and every other leaf below for a total of four sampled leaves per plant. These leaves were numbered sequentially from 1 (most apical) to 4 (most basal). We used four replicate plants per combination of P treatment and *Inv4m* genotype for a total of 64 tissue samples.

We took ten disc samples from the leaf tips with a tissue puncher and immediately froze the tissue in 1.5 mL tubes with two steel beads precooled with liquid nitrogen and kept in dry ice until stored at -80°C. We extracted total RNA with the QIAGEN RNAeasy Plant Mini Kit RNA extraction kit following manufacturer procedures (QIAGEN 74904), and RNA samples were quantified in nanodrop and sent to the NCSU Core Genomics Laboratory for sequencing. Following QC in Bioanalyzer, Illumina libraries were prepared and sequenced in a lane of Novaseq according to manufacturer recommendations.

### Plant genotyping

We followed (Brouard and Bissonnette 2022) for GATK-based RNAseq genotyping of 15 plant samples represented by 60 leaf libraries. Briefly, Illumina short reads were mapped to the NAM5 Zea mays B73 genome (Hufford *et al*. 2021) using *STAR* (Dobin *et al*. 2013), then we marked duplicates in the resulting BAM alignments, split reads at intron exon junctions and recalibrated sequence quality per leaf library.

At this point, we used HaplotypeCaller for generating gvcfs per plant identified by field row id (*∼* 4 libraries per plant). We did joint sample genotyping afterward with *genotypeGVCFs*. Then we filtered for variant quality (*window 35, cluster, QD < 2*.*0, FS > 30*.*0, SOR > 3*.*0, MQ < 40*.*0*) for the genotypes and 50% marker completion for individuals. This resulted in 200000 markers with 85% complete data for 13 plants. Finally, we used TASSEL5 K Nearest Neighbour imputing, producing a matrix of 19668 markers at 99.84% completion. Shell scripts are available at the *Inv4m* github repository

### Differential gene expression analysis

We aligned reads to the maize Zm-B73-REFERENCE-NAM-5.0 genome using *kallisto* (Bray *et al*. 2016). The alternative transcript alignment was turned into counts per gene. We used *voom* to calculate variance weights according to gene expression levels and counts were converted to log_2_(CPM) (Ritchie *et al*. 2015). We made a multivariate multiple regression for gene expression using *limma*. For the log transformed expression *Y*_*ijrs*_, from leaf stage *s*, in plant *r*, under phosphorus treatment *i*, with genotype *j*, we have:

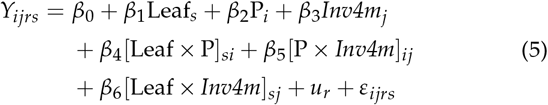

with plant level random effect and residuals:

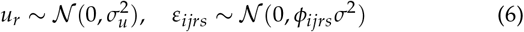

The term *u*_*r*_ represents a plant level random effect that accounts for non-independence among leaves sampled from the same individual. This approach correctly treats plants (n=4 per treatment*×* genotype) as biological replicates, with leaves as correlated sub-samples within each plant. The consensus within plant correlation was estimated using *duplicateCorrelation* in *limma* and incorporated into model fitting.

The leaf stage (Leaf_*s*_) was modeled as a centered numerical variable (*s∈* { 1, 2, 3, 4 } ), so the intercept represents expression at the average leaf stage. The precision weights *ϕ*_*ijrs*_ from the voom transformation capture heteroskedastic measurement error across samples.

We adjusted the p-values for the t-tests of the linear model coefficients as false discovery rates (FDR). Genes whose effect had FDR *<* 0.05 were deemed differentially expressed. For phosphorus treatment (*β*_2_) and *Inv4m* genotype (*β*_3_), genes with |log_2_(FC) | *>* 1.5 were considered strong DEGs. For the leaf stage effect (*β*_1_) and interaction terms, a threshold of |log_2_(FC) | *>* 0.5 was applied, corresponding to *>* 2.8-fold total change across the leaf gradient. R scripts and expression data are available at the *Inv4m* github repository.

### CDS sequence divergence analysis

To test whether expression differences in the introgression are explained by coding sequence divergence between haplotypes, we compared CDS percent identity with differential expression magnitude for genes in the shared introgression. CDS sequences for B73 (NAM v5) and the Mi21 NIL were aligned using LAST (Kiełbasa *et al*. 2011) as part of the microsynteny pipeline described above. For each B73 gene in the shared introgression with a reciprocal best hit in Mi21 NIL, we retained the highest scoring alignment and computed CDS divergence as *−*100 % identity. We restricted the analysis to CDS alignments because upstream regulatory regions in maize are frequently interrupted by polymorphic transposon insertions that make percent identity unreliable for non-coding sequences (Liang *et al*. 2022). Of the 512 expressed genes in the shared introgression, 254 had ortholog alignments and were included in the analysis (107 inside *Inv4m*, 147 in the flanking introgression). We fit five linear models predicting |log_2_ FC| as a function of CDS divergence, region (inside *Inv4m* vs flanking), their interaction, each term alone, and an intercept-only null, and selected the best model by BIC (Supplementary Table S7). Wilcoxon rank sum tests compared CDS divergence and log_2_ fold change distributions between regions.

### WGCNA module perturbation analysis

To assess how *Inv4m* disrupts gene coexpression structure, we performed weighted gene coexpression network analysis (WGCNA) (Langfelder and Horvath 2008) on the 465 DEGs (FDR *<* 0.05) using field expression data only. Signed weighted networks were constructed using soft thresholding power selected via `pickSoftThreshold()` to achieve approximate scale free topology (*R*^2^ *>* 0.8). Both CTRL and *Inv4m* conditions independently converged on *β* = 18, which was used as a shared power for all downstream analyses to ensure comparability.

Reference coexpression modules were defined from CTRL samples using the consensus network approach of (Shahan *et al*. 2018), which uses gene subsampling to identify relationships that are stable across parameter choices. The Leaf Gradient reference network was constructed using the same consensus pipeline applied to an independent gene set of 218 panicoid grass photosynthesis (*n* = 163) and senescence (*n* = 55) genes (Ojeda-Rivera *et al*. 2026), providing a topological control with comparable module count and gene number for the genotype label permutation test. In each of 1000 iterations, 80% of genes were randomly subsampled without replacement and WGCNA clustering was performed with *minModuleSize* uniformly sampled from 10 to 35 to avoid parameterization bias. Following (Shahan *et al*. 2018), the consensus coclustering matrix *C* was computed as:

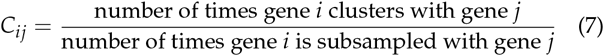

where *C*^*ij*^ *∈* [0, 1] represents the proportion of opportunities in which genes *i* and *j* were assigned to the same module. Unlike (Shahan *et al*. 2018) who randomized power transformation across iterations, we fixed soft power at *β* = 18 and varied only *minModuleSize* given our smaller gene set. Final CTRL reference modules were identified by hierarchical clustering on *C* (*deepSplit* = 4).

Module stability was assessed using 1000 independent bootstrap iterations (separate from the consensus iterations above), calculating the Largest Fragment Proportion (LFP), the proportion of each module’s genes captured by the largest bootstrap fragment. Module eigengenes were computed as the first principal component of each module’s expression matrix.

Network perturbation was quantified by comparing standard intramodular connectivity (*k*^*Within*^) between CTRL and *Inv4m* networks for genes assigned to CTRL consensus modules. For each genotype, signed weighted adjacency matrices were computed from the expression data using the shared soft thresholding power (*β* = 18). Intramodular connectivity was calculated using `intramodularConnectivity()` (Langfelder and Horvath 2008):

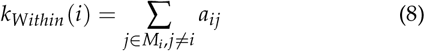

where *a*^*ij*^ is the signed adjacency between genes *i* and *j*, and *M*^*i*^ is the set of genes in the same module as gene *i*. Connectivity change was calculated as 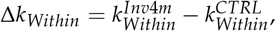, where negative values indicate connectivity loss in *Inv4m*. Module preservation was assessed using `modulePreservation()` (Langfelder *et al*. 2011) with 1000 label permutations, yielding *Z*^*summary*^ statistics (thresholds: *<* 2 = not preserved, 2 to 10 = moderate, *>* 10 = strong). Hub genes were identified as the top 5 genes per module ranked by *k*^*Within*^ in the CTRL network.

Gene Ontology enrichment analysis was performed for each module using *clusterProfiler* (Yu *et al*. 2012). Semantic redundancy was reduced using *rrvgo* (Sayols 2023) with a similarity threshold of 0.7, and the top Biological Process term per module was used to annotate the connectivity change boxplot.

### Trans coexpression network analysis

To identify genes regulated in *trans* by *Inv4m*, we constructed a bipartite coexpression network linking strong DEGs (|log_2_ FC |*>* 1.5) within the introgressed region (*cis* potential regulators) to strong DEGs located elsewhere in the genome (*trans* targets). Pearson correlation coefficients were calculated between all *cis*-*trans* gene pairs using log_2_(CPM) normalized expression values. Edges were retained if the correlation was significant after FDR correction (*q <* 0.05). Prior to network construction and to flowering time / plant height candidate selection (Table 1), we excluded *trans* DEGs carrying SNPs in linkage disequilibrium with the *Inv4m* tagging marker (Pearson correlation FDR *<* 0.005; see Section 3.1) because differential expression at these genes cannot be distinguished from cis effects of the linked SNPs. This filter removed 2 of 39 *trans* DEGs from the network (*Zm00001eb070810 / ychf* on chromosome 2 and *Zm00001eb123230 / tip4c* on chromosome 3); the same two genes were also excluded from Table 1. The filter was not applied to WGCNA module assignment, where the LD linked genes contribute to module composition but are not interpreted as candidate *Inv4m* regulators.

To distinguish *Inv4m* specific effects from flanking introgression effects, we applied a strongest correlated gene criterion: for each *trans* target, we identified the *cis* potential regulator with the highest absolute correlation. Targets were assigned to the *Inv4m* network only if their strongest regulator was located within the 15 Mb inversion proper, not in the flanking introgressed regions. This approach recovered 50 genes in the *Inv4m* trans regulatory network (22 upregulated, 28 downregulated).

To validate our coexpression edges against known regulatory relationships, we queried the MaizeNetome database (Feng *et al*. 2023), which provides genome wide gene coexpression networks derived from 7,603 maize RNA-seq samples. Because the MaizeNetome search interface limits batch queries, we retrieved edges in three batches using B73 NAM v4 gene identifiers: (1) left flanking region (upstream of *Inv4m*), (2) *Inv4m* proper (the 15 Mb inversion), and (3) right flanking region (downstream of *Inv4m*). Gene identifiers were converted from B73 NAM v4 to v5 using published correspondence tables (Hufford *et al*. 2021). Edges present in MaizeNetome were classified as “Reference” (representing conserved coexpression relationships detected across diverse maize tissues and genotypes), while edges absent from MaizeNetome were classified as “Dataset-specific” (condition specific to our field leaf samples). Of the 552 edges in the *Inv4m* trans regulatory network, 16 (2.9%) were Reference edges and 536 (97.1%) were dataset specific edges, indicating that the coexpression relationships detected in field grown V10–V12 leaves largely differ from those captured across the diverse tissue and genotype sampling of MaizeNetome.

Networks were visualized as minimum spanning trees (MST) constructed on standardized Euclidean distance derived from mutual information: *MI* = 0.5*×* log_2_(1/(1 *−r*^2^)). Edge polarity (concordant vs. discordant expression change) was indicated by arrow or bar symbols, respectively.

Gene Ontology enrichment analysis was performed on the trans network using *clusterProfiler* (Yu *et al*. 2012) with the B73 NAM v5 GO annotation from (Fattel *et al*. 2024). Gene sets were stratified by regulator location (flanking vs. *Inv4m*) and direction of differential expression (up vs. down). Semantic redundancy among enriched terms was reduced using *rrvgo* (Sayols 2023) with a similarity threshold of 0.7. Enrichment results were visualized as bubble plots showing the top five terms per gene set ranked by adjusted *p*-value.

### Filtering of Inv4m DEGs by phenotype association

As our data showed evidence of *Inv4m* accelerating flowering time and increasing plant height, we put together a list of candidate genes associated with these two phenotypes to tease out which DEGs were likely contributors to the observed *Inv4m* effect in these traits. For flowering time, we started with the list of 991 genes compiled by (Wang *et al*. 2021) and 62 genes from (Li *et al*. 2023).

Then we downloaded the maize data from the GWAS atlas (Liu *et al*. 2023) (*gwas_association_result_for_maize*.*txt*.*gz*) and selected genes that overlapped association SNPs for the Plant Phenotype and Trait Ontology term “days to flowering trait” *PPTO:0000155*. For this and the following candidate gene list, we considered that a gene overlapped an association SNP if the SNP was located within the 5 kb extended range of the gene model, i.e. as described in the gff gene annotation *±* 5 kb.

The final source of associations for flowering time was the phenotypic plasticity study in Tibbs-Cortes *et al*. (2024) from which we used 281 genes with significant GWAS SNPs in the columns *DTS_slope, DTS_intcp, DTA_slope, DTA_intcp*. For plant height, 27 genes from Liu *et al*. (2023), 1210 genes with GWAS Atlas associations for the term “plant height” *PPTO:0000126*; and 39 genes overlapping phenotypic plasticity association SNPs for *PH_slope* and *PH_intcp* (Tibbs-Cortes *et al*. 2024). The final nonredundant list consisted of a total of 2224 candidate genes for flowering time and 1272 candidates for plant height.

### Meristem clearing and size quantification

For vegetative meristem size quantification, maize seedlings were grown at the North Carolina State University Phytotron in a Percival Model LT-105 growth chamber (conditions: 29.4°C day/23.9°C night, relative humidity 50%, 16 hours light/8 hours dark, light intensity of 412 *µ* mol at plant height; soil type: 1:1 Sun Gro Propagation Growing Mix [Canadian Sphagnum peat moss 50-65%, vermiculite, dolomitic lime, 0.0001% silicon dioxide] : cement sand) in 24-well trays. Seedlings were watered daily in the morning and fertilized three times per week.

Two-weeks after planting, seedlings were cut at soil level and again 1cm above the soil cut. This 1cm tissue cassette of the shoot apex was cut longitudinally in the medial plane, in a midrib to margin orientation, by hand with a razor blade. Tissue was fixed in ice cold and fresh FAA [50% EtOH, 35% milliQ water, 10% formaldehyde (35%), 5% glacial acetic acid (v/v)] with a vacuum for 15 min. FAA was replaced, and tissues were place overnight at 4°C on a rocker.

For clearing, tissues were then removed from FAA and dehydrated through a graded ethanol series at room temperature for an hour each with gentle shaking: 50%, 70%, 85%, 95% EtOH (v/v)], followed by 1:1 95% EtOH and Methyl Salicylate, and finally, 100% Methyl Salicylate. Once in 100% Methyl Salicylate, samples were left to shake at room temperature overnight. Each shoot apex tissue cassette was placed on a microscope slide and covered with a coverslip.

Cleared shoot apices were imaged with differential internal contrast using a Leica DM4B microscope equipped with a DMC6200 digital camera. Each shoot apex image was measured using ImageJ v2.14.0. The scale was properly set for each image. Width was measured as a straight line at 0° anchored from the edge of P0. Height was measured from the highest point of the meristem tip to the width line at 90°. Surface area was measured using the polygon selection tool by taking into consideration the width line and marking the edge of the meristem dome.

Data were analyzed using ANOVA and plots were generated in R v.4.3.2 with the packages rstatix v.0.7.2, readxl v.1.4.3, ggplot2 v.3.5.1 and ggpubr v.0.6.0. SAM Welch *t*-tests in Figure 2D were one tailed (Inv4m *>* CTRL) and the BH-FDR pool was restricted to the three elongation related dimensions (height, *h*/*r, h*/*r*^2^).

## Acknowledgments

Fieldwork and mapping population development were supported by NSF-PGR award 1546719 This work is supported by the Research Capacity Fund (HATCH), project award nos. 7005660 and 7008935, from the U.S. Department of Agriculture’s National Institute of Food and Agriculture. This research was supported by the U.S. Department of Energy, Office of Science, Biological and Environmental Research program, Early Career Award Number DE-SC0021889. This research was supported by the United States Department of Agriculture (USDA) Agricultural Research Service (USDA-ARS). Additional support was provided by the Foundation for Food and Agriculture Research (FFAR) with support from the Grantham Foundation. This research was supported by the Science and Technologies for Phosphorus Sustainability (STEPS) Center, a National Science Foundation Science and Technology Center (CBET-2019435). Andi Kur designed and illustrated the cover graphic. We thank members of the Ruairidh Sawers and Rubén Rellán Álvarez labs for their contribution to population development and field work that made possible the research in this manuscript. We are also grateful to Jeff Ross-Ibarra and Rafael Guerrero for critical evaluation of the manuscript and feedback. We thank the Puerto Vallarta Winter Nursery crews who have helped generate introgression populations used in this manuscript. We thank the staff at Penn State’s Rock Springs and NC State’s Central Crops Research Stations for supporting field work shown in this paper. We especially want to acknowledge the indigenous people of the Americas and the ingenuity with which they domesticated and facilitated the spread and adaptation of maize throughout the continent. This work would not have been possible without the international maize research community and the willingness of so many colleagues to support the development of new research programs. Any opinions, findings, conclusions, or recommendations expressed in this publication are those of the author(s) and should not be construed to represent any official USDA, NSF, DOE, ARS or U.S. Government determination or policy.

## Supplementary material

**Figure S1.**
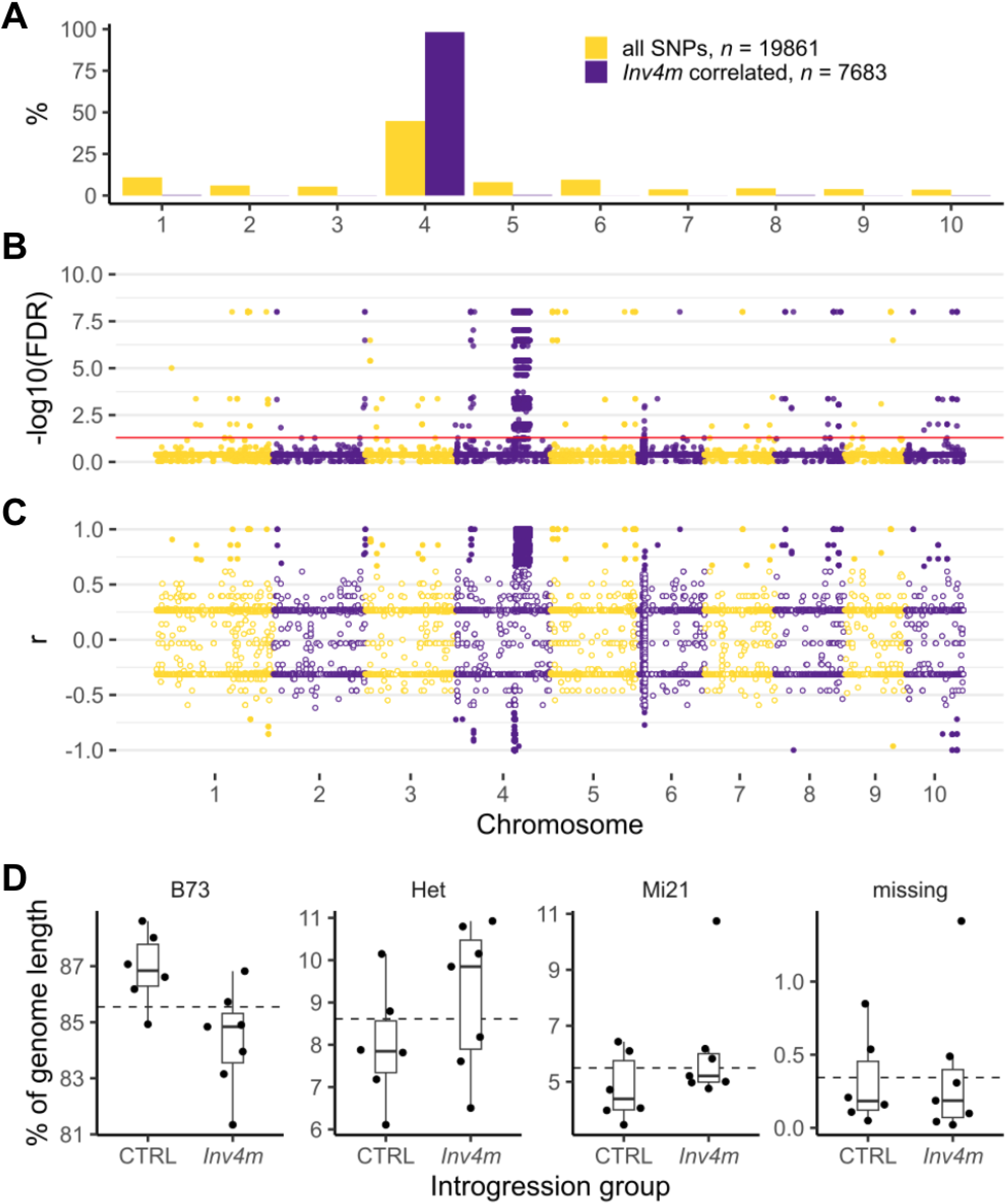
Chromosome Distribution of the SNPs in the RNAseq samples, their correlation with *Inv4m*, and genotype run length as percentage of the genome. **(A)** Chromosome 4 contains 44% of all genotyped SNPs and 98% of *Inv4m* correlated SNPs (n=13). **(B)** Manhattan plot for the significance (*t-test*) of the Pearson correlation of 19861 genotyped SNPs with PZE04175660223. Red line: *FDR* = 0.005 threshold. **(C)** Pearson correlation (*r*) of each SNP with the *Inv4m* tagging SNP PZE04175660223, open circles: non-significant SNPs. SNPs with significant correlation with *Inv4m* (full circles) are divergent highland introgressions. **(D)** NIL genotype run length as percentage of the genome. Each point represents a NIL, *Inv4m* plants are tagged by the highland allele of PZE04175660223, CTRL plants by the reference (B73) allele, dashed line is the overall mean for the 13 NILs.

**Figure S2.**
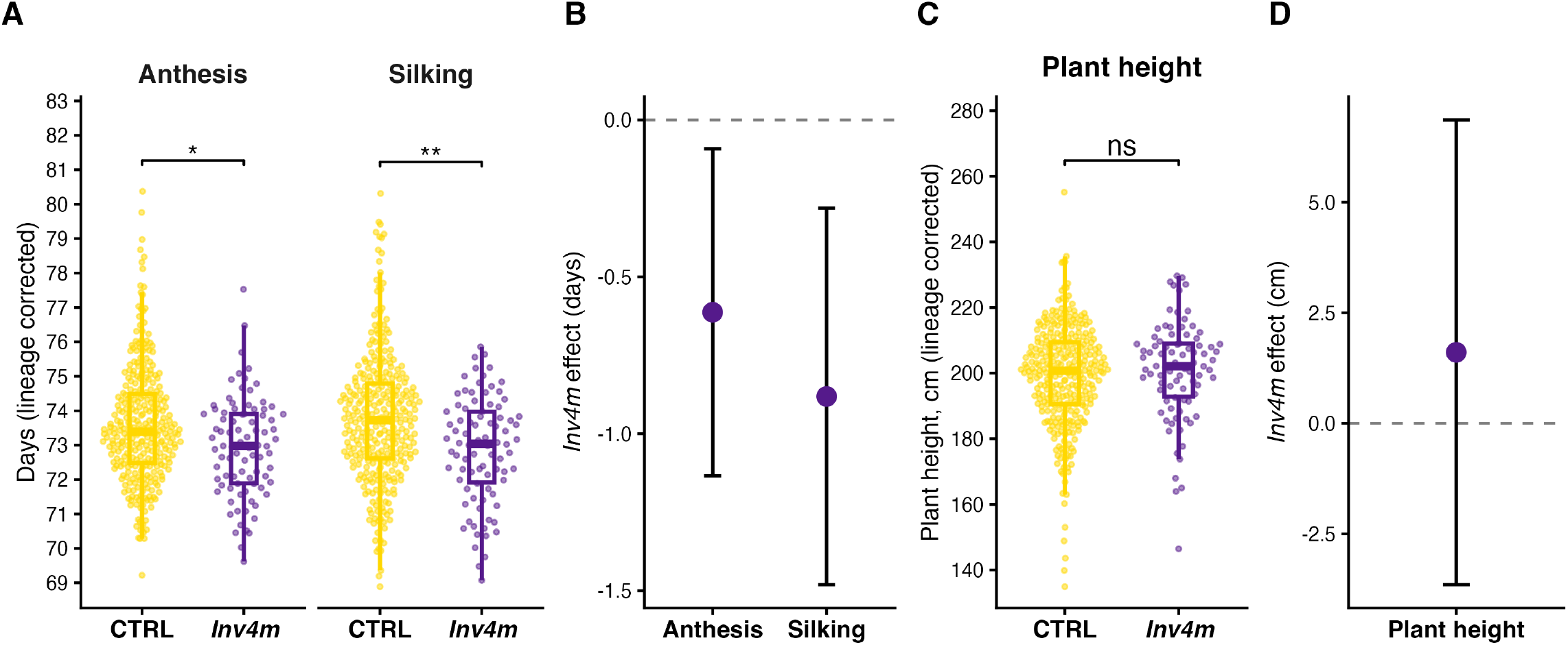
*Inv4m* effect on flowering and plant height in the ZeaL near isogenic line panel (NC2025). **(A)** Lineage corrected days to anthesis (DTA, left) and days to silking (DTS, right) for CTRL (*Inv4m* recurrent allele, B73 haplotype, gold) and *Inv4m* (donor allele, mexicana or huehuetenanguensis haplotype, purple). Each *Inv4m* arm shows the spread of NIL line means around its arm mean: ancestry and BC2 family random effect BLUPs from the linear mixed model were subtracted from the spatially corrected values, while NIL identity was retained so the visual spread reflects the line to line variation the model uses for inference. Stars indicate significance from the *Inv4m* fixed effect Satterthwaite *p* (Materials and Methods); ** *p <* 0.01, * *p <* 0.05. **(B)** Forest plot of the linear mixed model *Inv4m* fixed effect estimate (donor*−* recurrent) for DTA and DTS, with 95% Wald confidence intervals. **(C)** Lineage corrected plant height (PH); same conventions as A. **(D)** Forest plot of the *Inv4m* fixed effect estimate for PH; the donor effect is positive but not significant in this panel. *n* = 393 plots from 218 NILs across 35 BC1 families spanning five donor ancestry groups (Chalco, Durango, Mesa-Central, Nobogame, Huehuetenango). Donor lineages that do not carry the *Inv4m* mexicana haplotype (parviglumis, *Z. luxurians, Z. diploperennis*) are excluded by the both arms BC1 filter.

**Figure S3.**
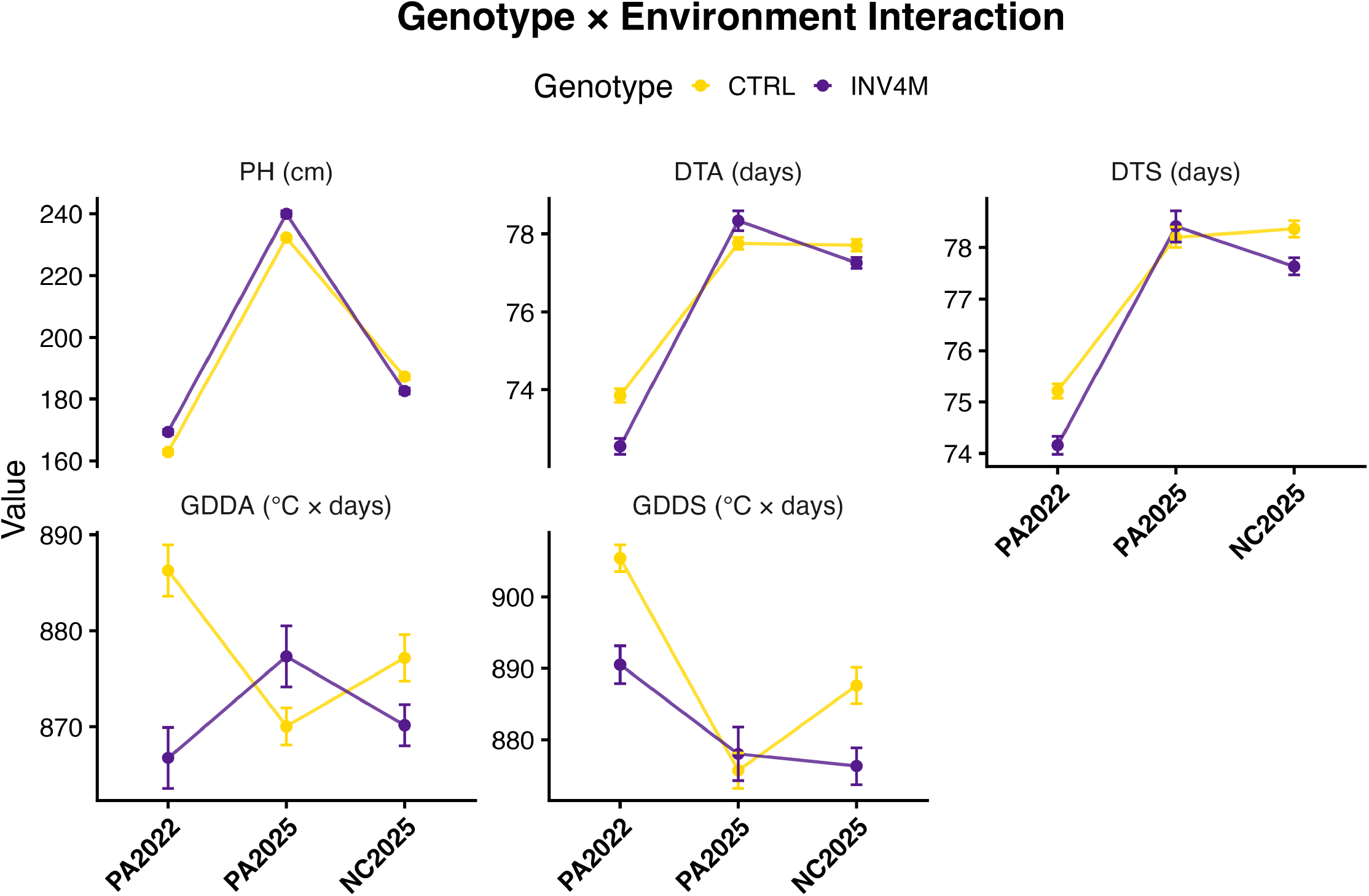
Genotype*×*environment interaction plots for the Mi21 donor background. Mean trait values (*±*SE) for CTRL (gold) and *Inv4m* (purple) genotypes across three environments: PA2022 and PA2025 (Pennsylvania) and NC2025 (North Carolina). Panels show days to anthesis (DTA), days to silking (DTS), plant height (PH), and thermal time equivalents (GDDA, GDDS). Non-parallel lines indicate genotype*×* environment interactions. Note the crossing interaction for plant height, where *Inv4m* is taller in Pennsylvania environments but shorter in North Carolina, consistent with environment dependent effects on growth.

**Figure S4.**
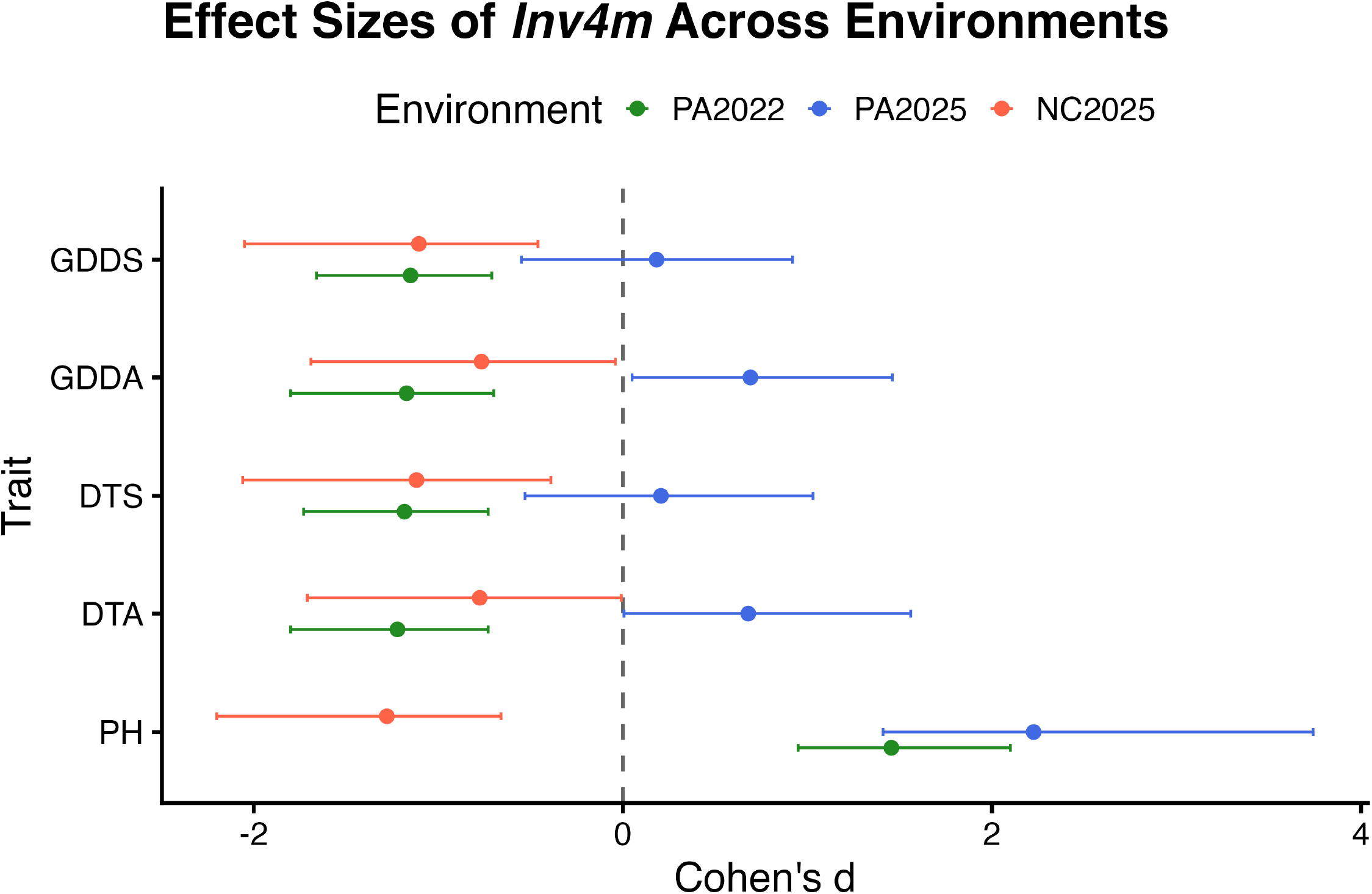
Effect sizes of *Inv4m* across environments. Forest plot showing Cohen’s *d* effect sizes for *Inv4m* genotype comparisons within each environment (PA2022, PA2025, NC2025) for plant height (PH), days to anthesis (DTA), days to silking (DTS), and thermal time equivalents (GDDA, GDDS). Error bars represent 95% confidence intervals. Note the reversal of plant height effects in NC2025 (North Carolina) compared to Pennsylvania environments.

**Figure S5.**
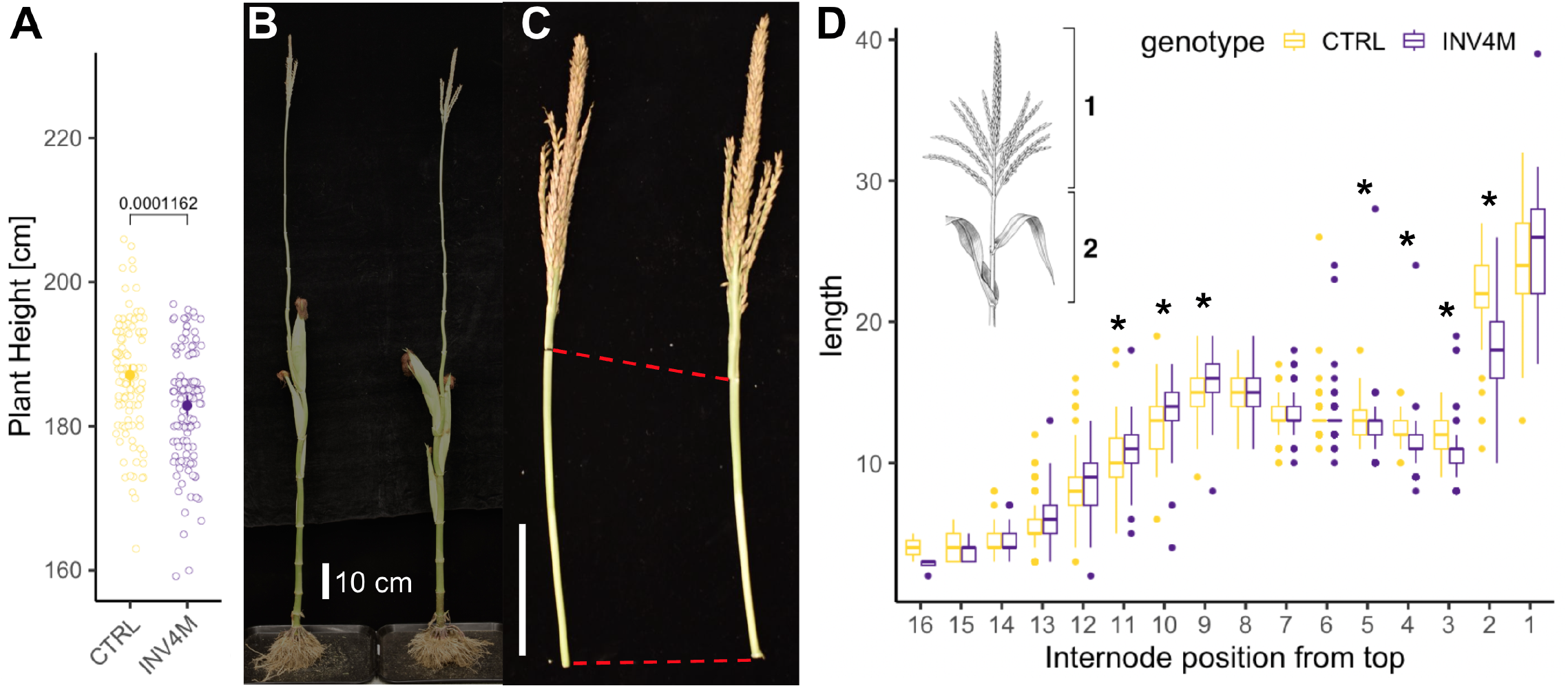
Internode length analysis in *Inv4m* and control plants. **(A)** Plant height comparison between CTRL and *Inv4m* in the Mi21 background (NC2025 field experiment). Boxplots show plant height (cm) with individual data points. P-value from linear mixed model. **(B)** Representative dissected plant showing whole plant architecture including root system and stem with visible node demarcations. **(C)** Detail of tassel and upper stem region. Scale bar = 10 cm. **(D)** Internode length profiles by position from top. Scale bar = 10 cm. Boxplots show internode length (cm) at each position for CTRL (gold, *n* = 112 plants) and *Inv4m* (purple, *n* = 116 plants). Asterisks indicate positions with significant differences (FDR *<* 0.05). Internode 2 shows the largest difference (*−* 4.12 cm, *−*18.5%; FDR = 3.7*×* 10^*−*18^) and accounts for 93% of the total height difference (Supplementary Table S5). *Inset*: Schematic illustration of maize plant architecture indicating internode numbering scheme from the tassel bearing node (position 1) downward.

**Figure S6.**
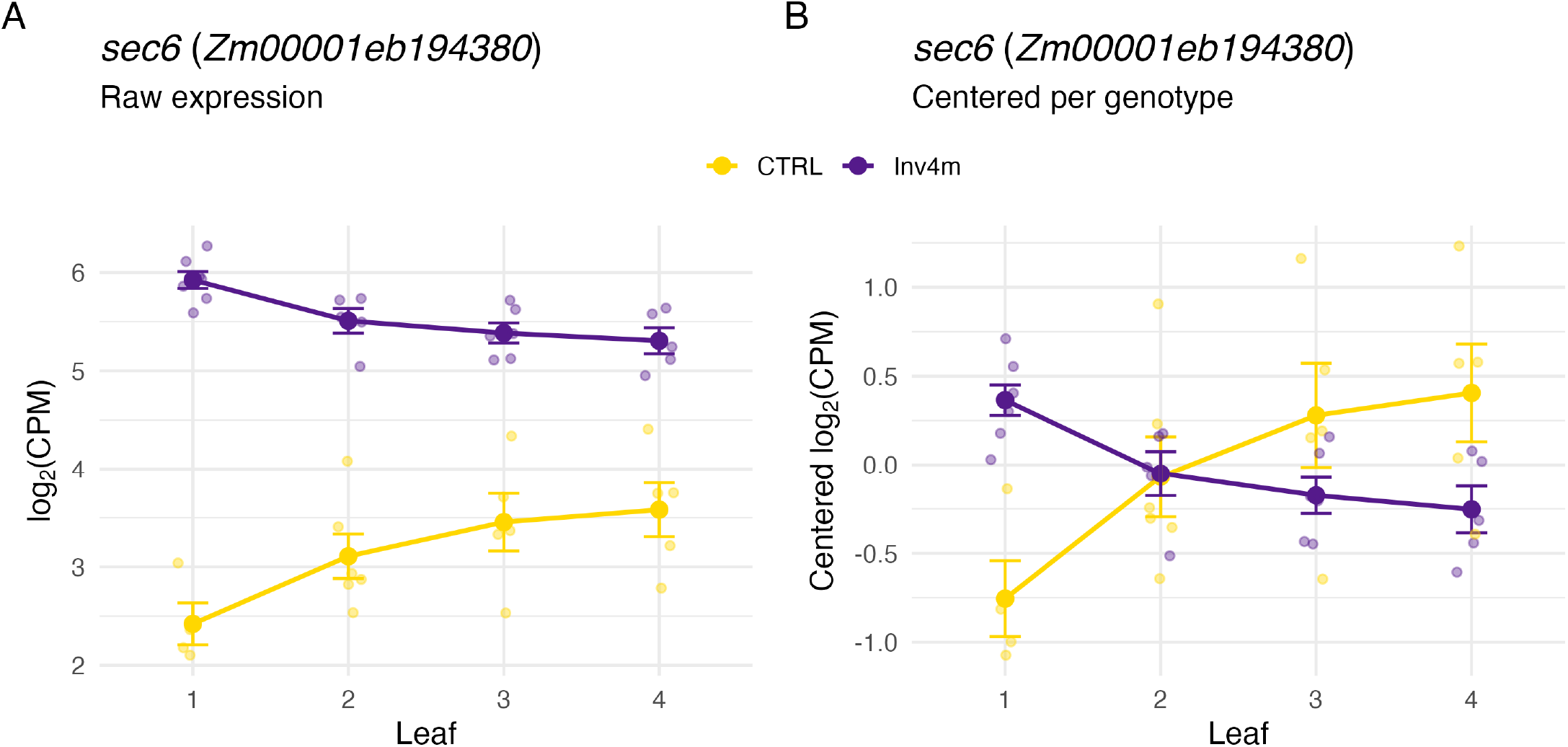
*sec6* expression across leaf positions in CTRL and *Inv4m*.

**Figure S7.**
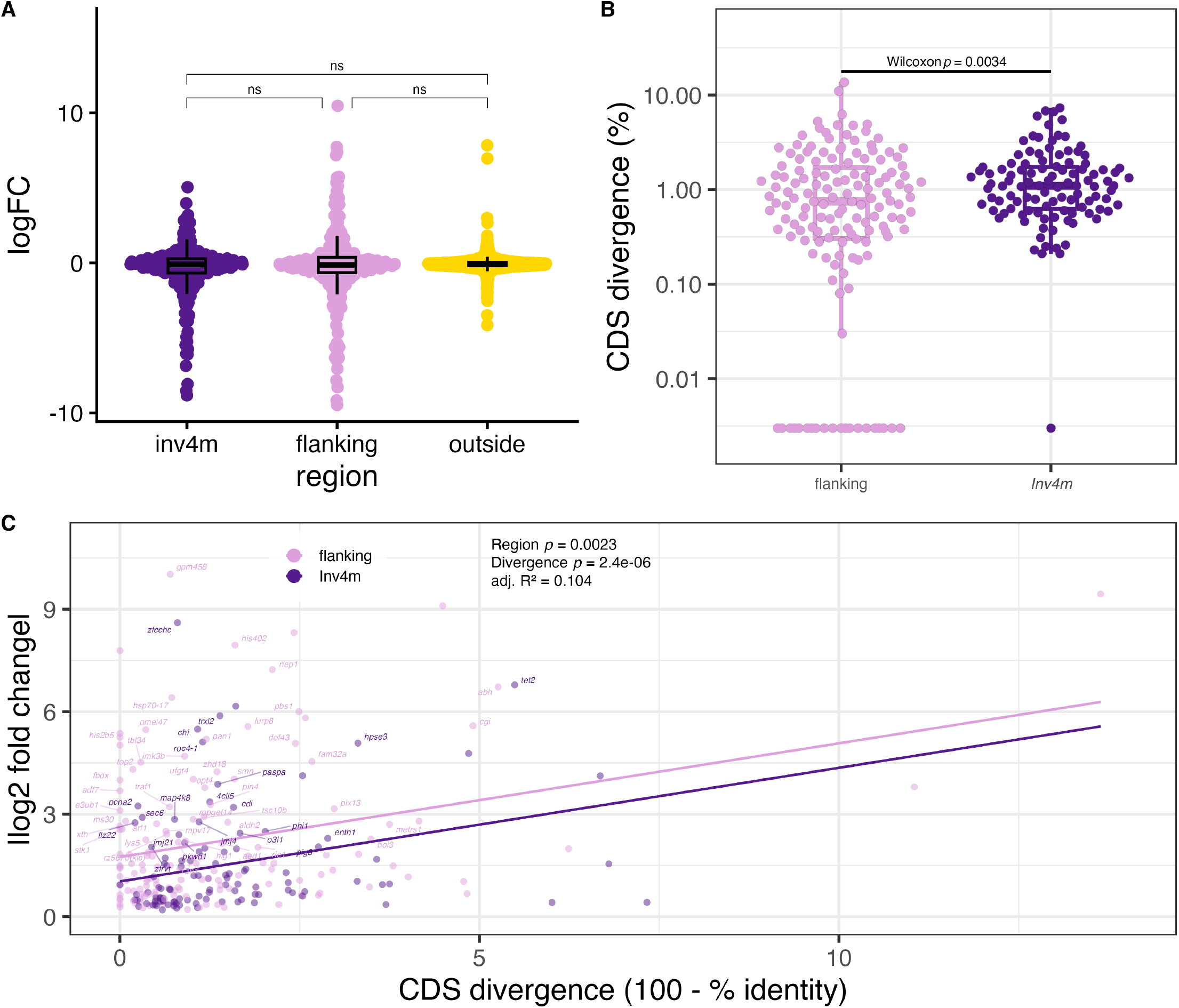
Expression change and CDS sequence divergence in the *Inv4m* introgression. **(A)** Distribution of log_2_ fold change for all expressed genes on chromosome 4, grouped by region (*Inv4m*, flanking introgression, outside). Wilcoxon rank sum tests show no significant directional bias between regions. **(B)** Distribution of CDS divergence (100 *−*% identity) for expressed genes inside the inversion (*Inv4m, n* = 107) and in the flanking introgression (flanking, *n* = 147). Points show individual genes (beeswarm); boxes show median and interquartile range. *y*-axis is log_10_-scaled. Wilcoxon rank sum *p* = 0.0034. **(C)** Absolute log_2_ fold change vs CDS divergence. Parallel lines show the additive model (shared slope for divergence effect, vertical offset for region effect). Region *p* = 0.0023; divergence *p* = 2.4e-06; adj. *R*^2^ = 0.104.

**Figure S8.**
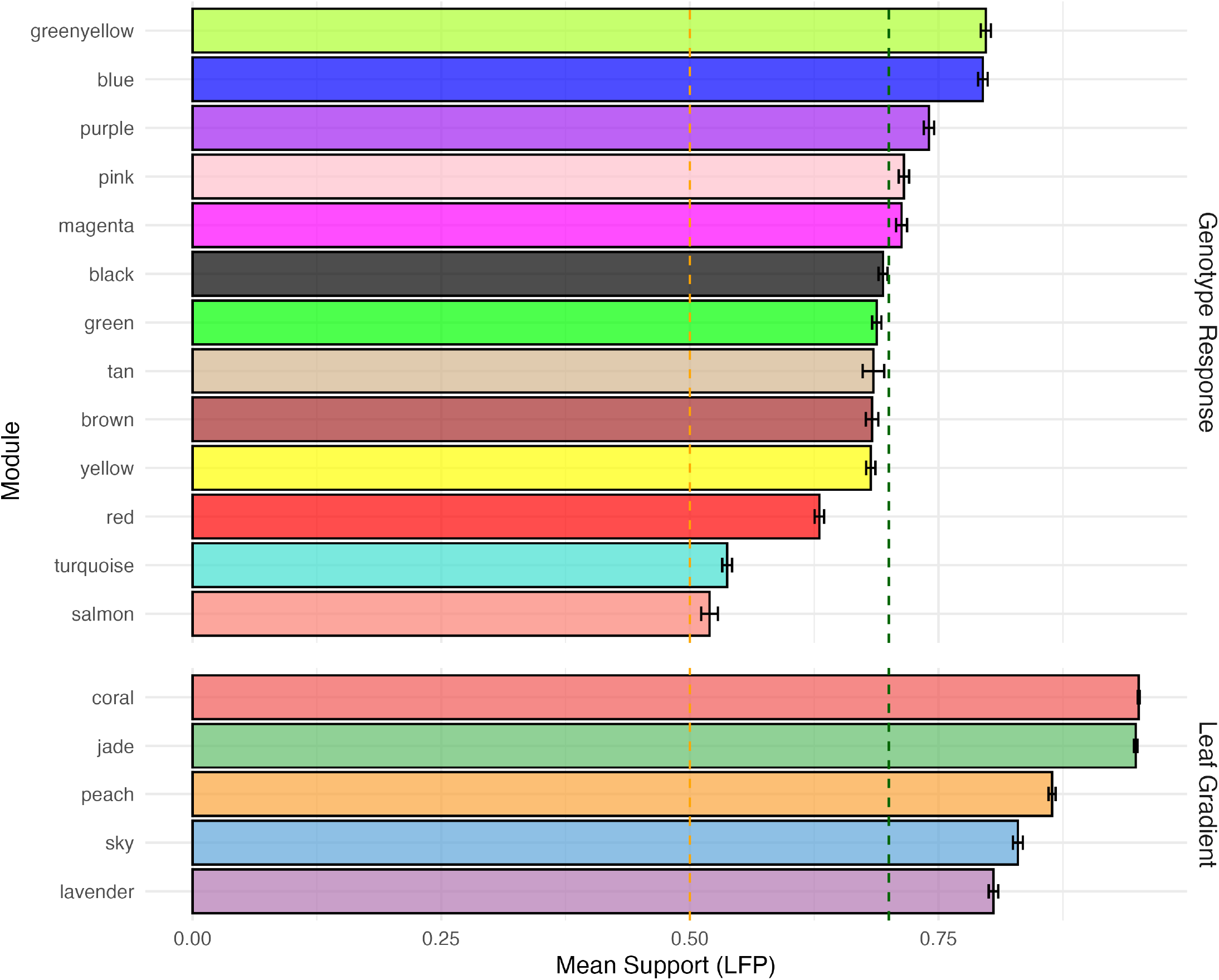
WGCNA consensus module bootstrap support for the Genotype Response and Leaf Gradient reference networks. Horizontal bars show mean bootstrap support (Largest Fragment Proportion, LFP) for each CTRL consensus module across 1000 iterations with 80% gene subsampling. Error bars indicate standard error of the mean. Dashed lines indicate stability thresholds: green (*≥*0.70, Stable) and orange (*≥*0.50, Moderate). *Top:* The 13 modules from the Genotype Response reference network, built from the 465 *Inv4m* DEGs (FDR *<* 0.05); bars are ordered by mean LFP. *Bottom:* The 5 modules from the Leaf Gradient reference network (coral, jade, peach, sky, lavender), which groups genes by their expression along the leaf developmental gradient and serves as an orthogonal control set of connectivity hubs. Most Genotype Response modules achieve moderate to stable support, and all Leaf Gradient modules reach stable support (*≥* 0.70), indicating robust module assignments in both CTRL reference networks.

**Figure S9.**
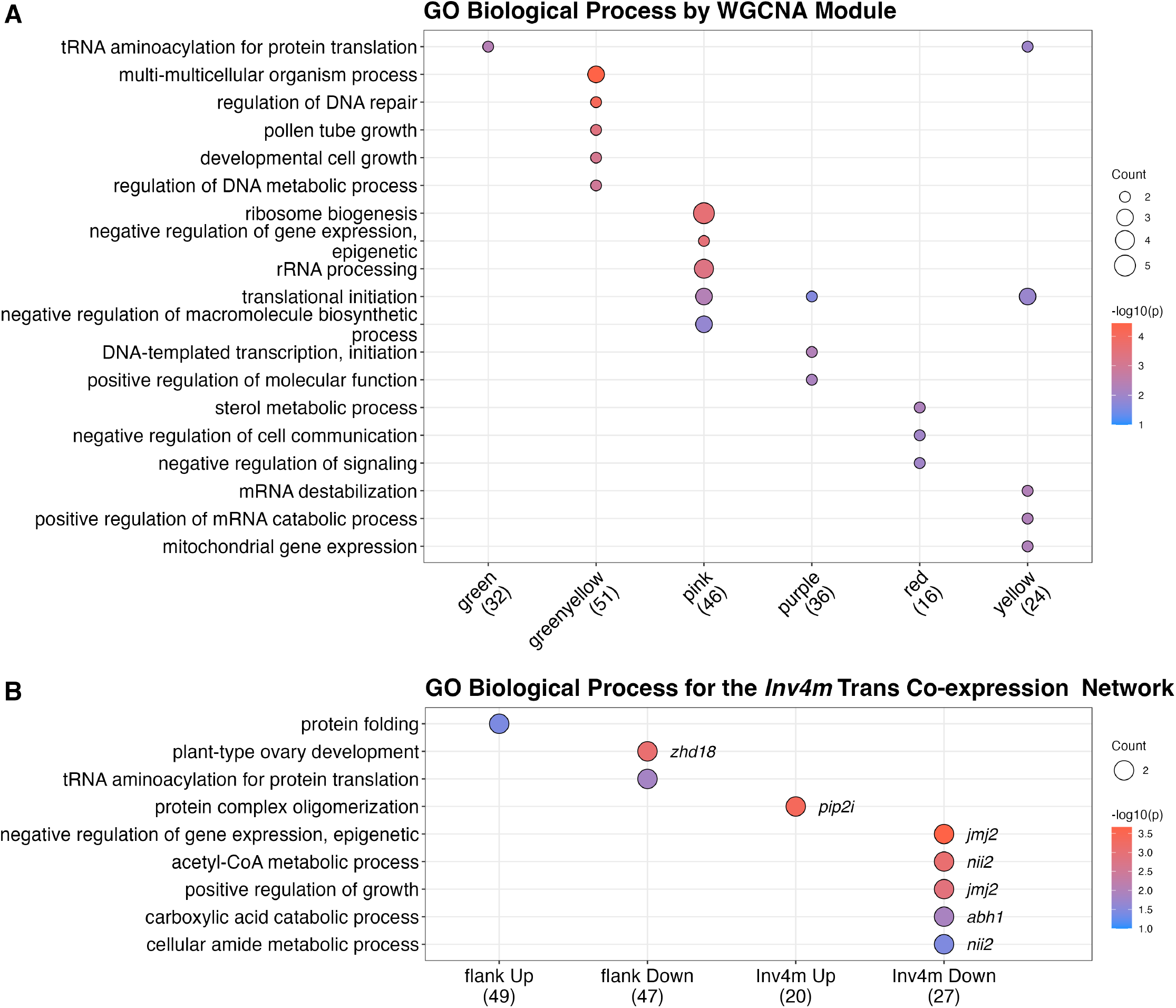
Gene Ontology enrichment of the WGCNA consensus modules and the *Inv4m* trans coexpression network. **(A)** Biological Process GO term enrichment for each CTRL WGCNA consensus module. Module labels include gene counts in parentheses. Notable enrichments include chromatin organization (blue module), translation (brown), and cellular response to stress (yellow). **(B)** GO enrichment of the *Inv4m* trans coexpression network from Fig 4C, stratified by gene location (flanking introgression vs. *Inv4m* proper) and regulation direction. Notable enrichments include negative regulation of gene expression (epigenetic) and positive regulation of growth among *Inv4m* downregulated genes. In both panels, bubble size indicates gene count and color indicates enrichment significance (*−* log_10_(*p*)); GO terms were reduced using rrvgo semantic similarity to eliminate redundancy.

**Figure S10.**
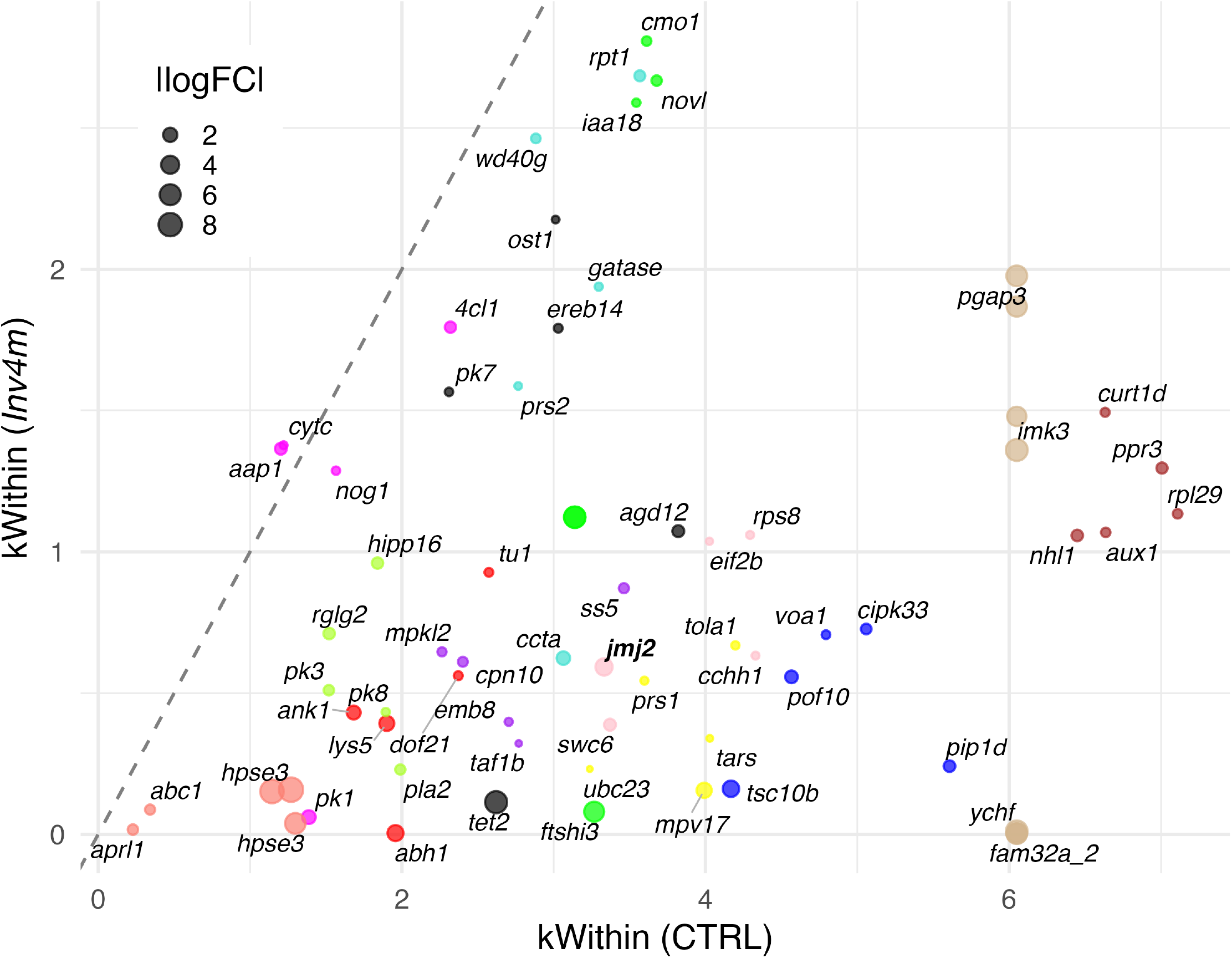
Intramodular hub gene connectivity in CTRL vs. *Inv4m*. Each point shows one of the top five hub genes (by CTRL *k*_*Within*_) in each WGCNA consensus module; point color corresponds to module identity as in Fig 4. The x-axis is intramodular connectivity in the CTRL consensus network and the y-axis is connectivity in the *Inv4m* consensus network; points below the dashed identity line have lost connectivity in the *Inv4m* karyotype. Point size encodes |log_2_ FC| from the genotype DEG analysis. Gene labels show curated names where available; *jmj2* is shown in bold italic. This figure complements Fig 4B, which summarizes module level connectivity loss as a permutation based *z*-score.

**Figure S11.**
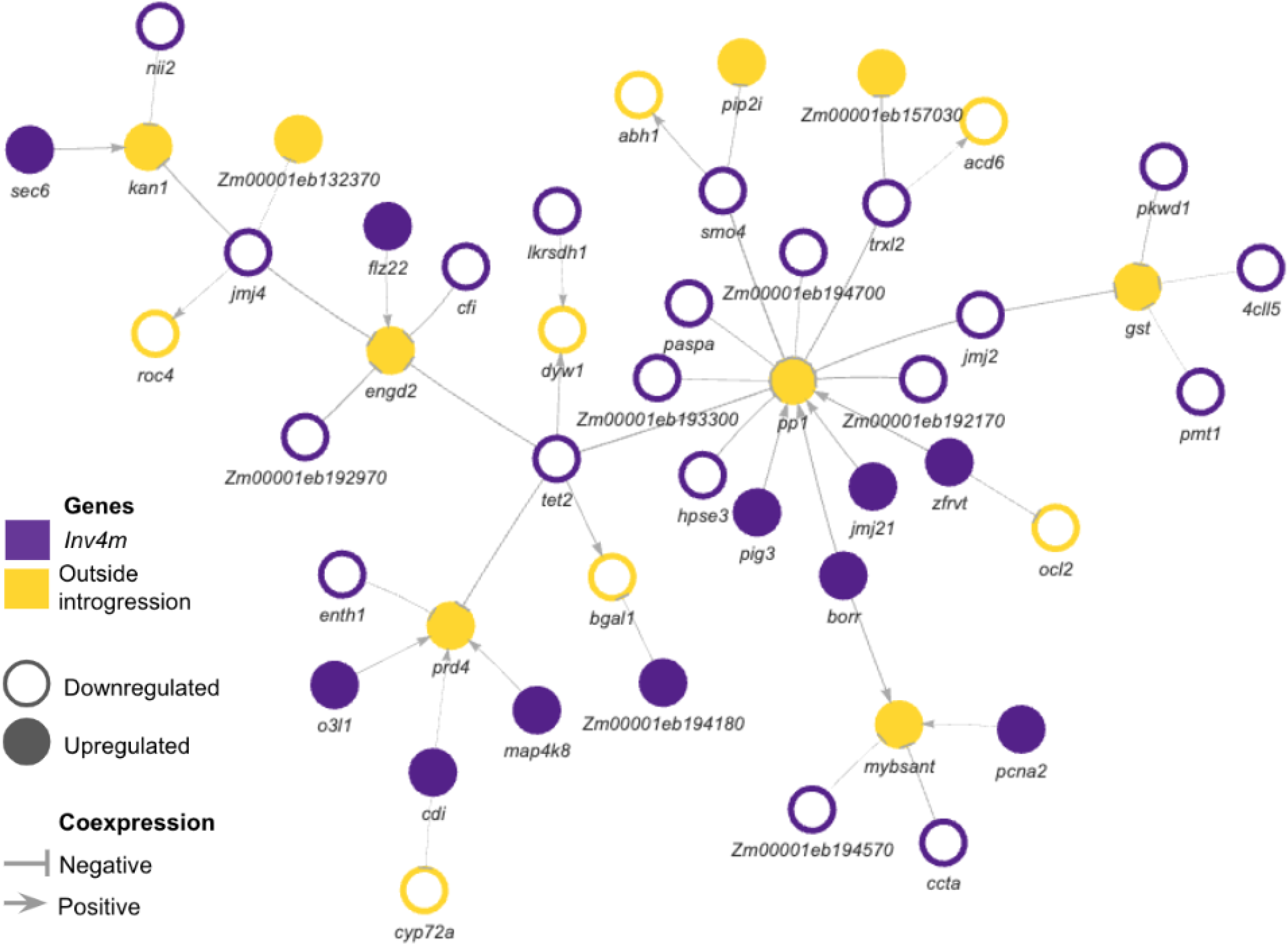
Full *Inv4m* dataset specific trans coexpression network (Bonferroni *<* 0.001). Minimum spanning tree of all dataset specific coexpression edges not present in the MaizeNetome reference, filtered to Bonferroni-adjusted *p <* 0.001 across the cis *×* trans correlation test pool (*m≈* 1,095 tests, *n* = 43 samples). Node fill indicates genomic location: purple (*Inv4m* region), pink (flanking introgression), gold (outside inversion). Colored fill indicates upregulation in *Inv4m*; white fill with colored border indicates downregulation. Edge width is proportional to mutual information. Gene labels are shown in italics. This figure shows the complete novel network; the *jmj4* neighborhood subgraph is detailed in Fig 4C.

**Figure S12.**
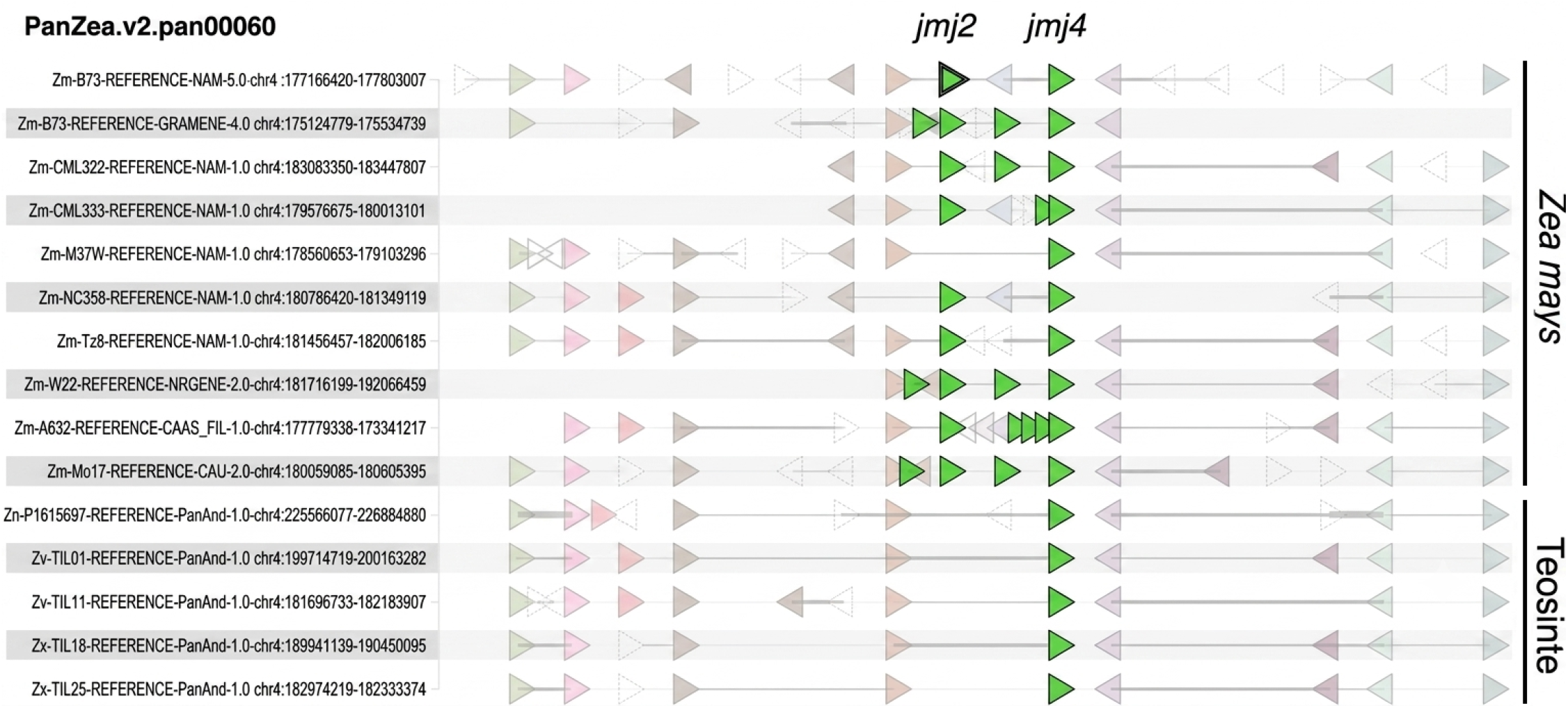
Microsynteny of the *jmj2*/ *jmj4* cluster across *Zea mays* and teosinte genomes. PanZea v2 pangene track (PanZea.v2.pan00060) of the genomic neighborhood around *jmj2* and *jmj4* in nine *Zea mays* NAM reference assemblies (top) and five teosinte assemblies (bottom; *parviglumis, mexicana*, and *Z. nicaraguensis* lines). Arrows represent gene models colored by pangene ortholog group; green arrows mark the *jmj2*/ *jmj4* tandem array. The *Zea mays* assemblies carry the multi copy tandem array described in Fig 5B, whereas the teosinte assemblies retain a reduced or single copy configuration, consistent with a lineage specific expansion in *Z. mays*. Interactive view: https://gcv.maizegdb.org/gene;maize=Zm00001eb191810?sources=maize&neighbors=10.

**Table S1.** Distribution of *Inv4m* DEGs in the Maize Genome.

| Location of genes | Total<br>Annotated | Total<br>Expressed | Fraction<br>Expressed | DEGs | Strong DEGs |
| --- | --- | --- | --- | --- | --- |
| | | | | $FDR < 0.05$ | $FDR < 0.05$<br>$\vert \log_2FC\vert > 1.5$ |
| Chromosomes 1-10 | 43459 | 24011 | 0.55 | 465 | 155 |
| Outside Introgression | 42360 | 23403 | 0.55 | 193 | 39 |
| Shared Introgression | 1099 | 608 | 0.55 | 272 | 116 |
| *Inv4m* | 432 | 253 | 0.59 | 114 | 40 |
| Flanking Introgression | 667 | 355 | 0.53 | 158 | 76 |

| Comparison | DEGs Enrichment |  |  | Strong DEGs Enrichment |  |  |
| --- | --- | --- | --- | --- | --- | --- |
| | Odds<br>Ratio | Fisher's<br>Exact test<br>$p$-value | Confidence<br>Interval | Odds<br>Ratio | Fisher's<br>Exact test<br>$p$-value | Confidence<br>Interval |
| Shared introgression vs Outside | 97.1 | $< 10^{-200}$ | [70,135] | 141 | $< 10^{-100}$ | [95,215] |
| Flanking vs Outside | 96.5 | $< 10^{-150}$ | [68,135] | 163 | $< 10^{-80}$ | [100,275] |
| ***Inv4m* vs Outside** | **98.7** | $< 10^{-100}$ | [65,150] | **112** | $< 10^{-45}$ | [65,200] |
| *Inv4m* vs Flanking | 1.02 | 0.95 | [0.7,1.5] | 0.69 | 0.25 | [0.4,1.2] |

**Table S2.** *Inv4m* delimitation and breakpoints.

|  | Start | End | Length | Start gene | End gene |
| --- | --- | --- | --- | --- | --- |
| TIL18 | 180365316 | 193570652 | 13205336 | *Zx00002aa015554* | *Zx00002aa015905* |
| PT | 186925484 | 173486186 | 13439298 | *Zm00109aa017629* | *Zm00109aa018009* |
| Mi21 NIL | 156752807 | 167620500 | 10867693 | *Zm00001eb194800* | *Zm00001eb190470* |
| B73 | 172883881 | 188131462 | 15247580 | *Zm00001eb190470* | *Zm00001eb194800* |

***Inv4m* Breakpoints**
| Position | Upstream Segment |  |  | Downstream Segment |  |  |
| --- | --- | --- | --- | --- | --- | --- |
|  | Start | End | Span | Start | End | Span |
| TIL18 | 180269950 | 180365316 | 95366 | 193570651 | 193734606 | 163955 |
| PT | 173369064 | 173486186 | 117122 | 186925483 | 187092654 | 167171 |
| Mi21 NIL | 156560061 | 156750619 | 190558 | 167621241 | 167838764 | 217523 |
| B73 | 172561330 | 172883881 | 322551 | 188131461 | 188220418 | 88957 |

| Genes | Upstream bounding |  | Downstream bounding |  |
| --- | --- | --- | --- | --- |
| TIL18 | *Zx00002aa015905* | *Zx00002aa015906* | *Zx00002aa015906* | *Zx00002aa015554* |
| PT | *Zm00109aa017627* | *Zm00109aa017629* | *Zx00002aa015905* | *Zm00109aa017629* |
| Mi21 NIL | *Zm00001eb190450* | *Zm00001eb194800* | *Zm00001eb190470* | *Zm00001eb194820* |
| B73 | *Zm00001eb190450* | *Zm00001eb190470* | *Zm00109aa017627* | *Zm00001eb194820* |

**Table S3.** *Inv4m* breakpoint knob repeats. Distribution of 180 bp knob repeats in the breakpoint segments and internal to the inversion.

| Upstream Breakpoint Segment |  |  |  |  |  |  |  |
| --- | --- | --- | --- | --- | --- | --- | --- |
|  | Repeat<br>count | Repeat<br>match<br>length | Start | End | Span | % Knob<br>Span | % Knob |
| TIL18 | 44 | 7638 | 180271106 | 180292951 | 21845 | 23% | 35% |
| PT | 44 | 7638 | 173370220 | 173392066 | 21846 | 19% | 35% |
| Mi21 NIL | 210 | 36316 | 167622041 | 167760234 | 138193 | 64% | 26% |
| B73 | 352 | 59580 | 172561959 | 172882309 | 320350 | 99% | 19% |
| *Inv4m* Internal Knob Repeat |  |  |  |  |  |  |  |
|  | Repeat<br>count | Repeat<br>match<br>length | Start | End | Span |  |  |
| TIL18 | 406 | 69913 | 183816987 | 184072303 | 255316 |  |  |
| PT | 354 | 61103 | 177250166 | 177516302 | 266136 |  |  |
| Mi21 NIL | 330 | 57545 | 160494941 | 160755992 | 261051 |  |  |
| B73 | 650 | 111841 | 184152738 | 184525111 | 372373 |  |  |
| Downstream Breakpoint Segment |  |  |  |  |  |  |  |
|  | Repeat<br>count | Repeat<br>match<br>length | Start | End | Span | % Knob<br>Span | % Knob |
| TIL18 | 154 | 25847 | 193572198 | 193671081 | 98883 | 59% | 26% |
| PT | 130 | 21466 | 186927027 | 187051351 | 124324 | 74% | 17% |
| Mi21 NIL | 57 | 9865 | 156560208 | 156615508 | 55300 | 29% | 18% |
| B73 | 26 | 4367 | 188149304 | 188219984 | 70680 | 79% | 6% |

**Table S4.** Genotype*×* Environment interaction statistics. Two-way ANOVA results for *Inv4m* effects across three location-year combinations (PA2022, PA2025, NC2025) using the Mi21 donor background. Interaction *p*-values and effect sizes 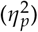 are shown for each phenotype.

| Trait | G × E *p*-value | G × E $\eta_p^2$ | Genotype *p* | Environment *p* | G × E sig. | G sig. |
| --- | --- | --- | --- | --- | --- | --- |
| PH | $8.7 \times 10^{-10}$ | 0.29 | $1.5 \times 10^{-7}$ | $3.2 \times 10^{-106}$ | *** | *** |
| DTA | $2.9 \times 10^{-5}$ | 0.16 | $2.0 \times 10^{-4}$ | $2.4 \times 10^{-54}$ | *** | *** |
| DTS | $4.6 \times 10^{-3}$ | 0.08 | $4.4 \times 10^{-5}$ | $4.1 \times 10^{-44}$ | ** | *** |
| GDDA | $7.1 \times 10^{-5}$ | 0.14 | $1.1 \times 10^{-4}$ | 0.51 | *** | *** |
| GDDS | $6.1 \times 10^{-3}$ | 0.08 | $1.9 \times 10^{-5}$ | $1.7 \times 10^{-13}$ | ** | *** |

**Table S5.** Internode length differences between *Inv4m* and control plants by position. T-tests comparing internode length at each position from the tassel (position 1) downward. FDR correction applied across all 16 positions. Internode 2 (immediately below the tassel) accounts for 93% of the total height difference. Mi21 donor background, NC2025 field experiment (*n* = 112 CTRL, *n* = 116 *Inv4m* plants).

| Position | CTRL (cm) | *Inv4m* (cm) | Diff (cm) | % Change | FDR | Sig. |
| --- | --- | --- | --- | --- | --- | --- |
| 1 | 24.19 | 25.09 | +0.91 | +3.8 | 0.150 | ns |
| 2 | **22.27** | **18.15** | **-4.12** | **-18.5** | **$3.7 \times 10^{-18}$** | **\*\*\*\*** |
| 3 | 11.86 | 10.73 | -1.12 | -9.5 | $3.0 \times 10^{-6}$ | **\*\*\*\*** |
| 4 | 12.38 | 11.28 | -1.09 | -8.8 | $1.4 \times 10^{-11}$ | **\*\*\*\*** |
| 5 | 12.98 | 12.40 | -0.59 | -4.5 | $3.8 \times 10^{-4}$ | **\*\*\*** |
| 6 | 13.17 | 13.02 | -0.15 | -1.2 | 0.245 | ns |
| 7 | 13.13 | 13.19 | +0.06 | +0.5 | 0.713 | ns |
| 8 | 15.22 | 15.17 | -0.05 | -0.3 | 0.816 | ns |
| 9 | 15.11 | 15.86 | +0.75 | +5.0 | $9.4 \times 10^{-4}$ | **\*\*\*** |
| 10 | 13.09 | 14.12 | +1.03 | +7.9 | $9.0 \times 10^{-5}$ | **\*\*\*\*** |
| 11 | 10.68 | 11.43 | +0.75 | +7.0 | 0.004 | **\*\*** |
| 12 | 8.62 | 9.00 | +0.38 | +4.5 | 0.240 | ns |
| 13 | 5.82 | 6.16 | +0.34 | +5.9 | 0.238 | ns |
| 14 | 4.43 | 4.51 | +0.08 | +1.9 | 0.713 | ns |
| 15 | 4.17 | 3.81 | -0.36 | -8.7 | 0.245 | ns |
| 16 | 4.00 | 2.75 | -1.25 | -31.3 | 0.238 | ns |

**Table S6.** Effect of *Inv4m* on gene expression. Selected DEGs from the plant blocking limma model (FDR *<* 0.05, | log_2_ FC| *>* 1.5). Bottom rows show significant Leaf*×Inv4m* interactions.

| Gene ID | Label | $-\log_{10}(FDR)$ | $\log_2 FC$ | Description |
| --- | --- | --- | --- | --- |
| ***Inv4m*: Downregulated** |  |  |  |  |
| Zm00001eb191790 | *jmj2* | 21.9 | -3.49 | JUMONJI-transcription factor 2 |
| Zm00001eb191820 | *jmj4* | 19.7 | -2.84 | JUMONJI-transcription factor 4 |
| Zm00001eb189910 | *metrs1* | 17.5 | -2.86 | Methionyl-tRNA synthetase |
| Zm00001eb192850 | *smo4* | 17.5 | -1.98 | Ribosome biogenesis protein NOP53 |
| Zm00001eb192630 | *ccta* | 15.8 | -1.69 | CCT-alpha |
| Zm00001eb188070 | *arf1* | 15.8 | -2.78 | ADP-ribosylation factor |
| Zm00001eb188750 | *hsp70-17* | 14.2 | -6.48 | Heat shock 70 kDa protein 17 |
| Zm00001eb190240 |  | 14.2 | -9.16 |  |
| Zm00001eb188240 | *ropgef14* | 13.6 | -2.98 | Rop guanine nucleotide exchange facto... |
| Zm00001eb193260 | *trxl2* | 13.2 | -6.10 | Thioredoxin-like 3-1, chloroplastic |
| Zm00001eb190350 |  | 13.0 | -1.83 | Ankyrin repeat-containing protein |
| ***Inv4m*: Upregulated** |  |  |  |  |
| Zm00001eb264460 | *ychf* | 16.9 | +6.50 |  |
| Zm00001eb194380 | *sec6* | 16.5 | +2.87 | Exocyst complex component Sec6 |
| Zm00001eb190670 | *zfrvt* | 15.6 | +1.97 |  |
| Zm00001eb191990 | *pig3* | 15.0 | +1.97 | Quinone oxidoreductase PIG3 |
| Zm00001eb241410 | *etfa* | 14.0 | +4.46 | Electron transfer flavoprotein subuni... |
| Zm00001eb213070 | *gbp1* | 13.7 | +8.96 | GTP-binding protein related |
| Zm00001eb195370 |  | 13.6 | +1.63 |  |
| Zm00001eb041890 | *kan1* | 13.6 | +1.68 | KANADI1 |
| Zm00001eb189920 | *aldh2* | 13.6 | +2.70 | aldehyde dehydrogenase2 |
| Zm00001eb190750 | *jmj21* | 13.5 | +2.28 | JUMONJI-transcription factor 21 |
| Zm00001eb189580 |  | 13.2 | +7.38 |  |
| Zm00001eb182850 | *mybsant* | 12.4 | +6.96 |  |
| Zm00001eb112470 | *engd2* | 12.3 | +6.14 |  |
| Zm00001eb187280 | *gpm458* | 11.2 | +10.47 | Oil body associated protein like |
| **Leaf × *Inv4m*** |  |  |  |  |
| Zm00001eb194380 | *sec6* | 3.1 | -0.55 | Exocyst complex component Sec6 |

**Table S7.** Model comparison for the relationship between CDS divergence and expression change in the *Inv4m* introgression. Five linear models predicting |log_2_FC| were compared by BIC and AIC. The additive model (CDS divergence + region) provided the best fit, with the interaction term adding no improvement. Region distinguishes genes inside *Inv4m* from the flanking introgression.

| Model | df | AIC | BIC | adj. $R^2$ |
| --- | --- | --- | --- | --- |
| CDS divergence + region | 3 | 1032.2 | 1046.3 | 0.1037 |
| CDS divergence | 2 | 1039.6 | 1050.2 | 0.0734 |
| CDS divergence × region | 4 | 1033.2 | 1050.9 | 0.1035 |
| Region | 2 | 1052.8 | 1063.4 | 0.0242 |
| Null (intercept only) | 1 | 1058 | 1065.1 | 0 |

**Table S8.** WGCNA module preservation statistics for the Genotype Response (Inv4m DEG) reference network. Modules sorted by FDR from the genotype label permutation test used in Fig 4B (1000 shuffles of the CTRL/*Inv4m* labels). Δ*k*_*Within*_ is the mean intramodular connectivity change (*Inv4m−* CTRL); *Z*_Δ*k*_ is that observed value expressed as a *z*-score relative to the permutation null. *Z*_*density*_, *Z*_*connectivity*_, and *Z*_*summary*_ from `modulePreservation()` with 1000 permutations (Langfelder *et al*. 2011): *Z*_*summary*_ *<* 2 = not preserved, 2–10 = moderate, *>* 10 = strong. *Z*_*connectivity*_ quantifies rank preservation of intramodular connectivity; *Z*_*density*_ quantifies preservation of average connectivity strength. LFP = Largest Fragment Proportion (bootstrap support in CTRL).

| Module | Genes | $Z_{density}$ | $Z_{connectivity}$ | $Z_{summary}$ | $\Delta k_{Within}$ | $Z_{\Delta k}$ | FDR | LFP |
| --- | --- | --- | --- | --- | --- | --- | --- | --- |
| brown | 46 | 3.9 | 3.8 | 3.8 | -2.36 | -3.74 | $< 10^{-3}$ | 0.68 |
| tan | 12 | 2.5 | 2.0 | 2.2 | -3.01 | -3.03 | 0.010 | 0.68 |
| purple | 22 | 4.4 | 3.3 | 3.8 | -1.11 | -2.46 | 0.034 | 0.74 |
| blue | 51 | 1.9 | 4.4 | 3.1 | -1.64 | -2.37 | 0.034 | 0.79 |
| pink | 30 | 5.1 | 6.4 | 5.8 | -1.36 | -2.28 | 0.034 | 0.72 |
| yellow | 36 | 2.6 | 4.4 | 3.5 | -1.47 | -1.89 | 0.085 | 0.68 |
| salmon | 11 | 0.7 | 1.7 | 1.2 | -0.39 | -1.35 | 0.220 | 0.52 |
| greenyellow | 16 | 3.5 | 2.4 | 3.0 | -0.63 | -1.26 | 0.220 | 0.80 |
| red | 34 | 4.4 | 2.2 | 3.3 | -0.51 | -1.13 | 0.220 | 0.63 |
| black | 32 | 5.1 | 3.9 | 4.5 | -0.80 | -1.12 | 0.220 | 0.69 |
| green | 36 | 7.8 | 2.2 | 5.0 | -0.61 | -0.82 | 0.264 | 0.69 |
| magenta | 24 | 7.7 | 2.9 | 5.3 | -0.14 | -0.36 | 0.383 | 0.71 |
| turquoise | 55 | 6.5 | 5.6 | 6.1 | -0.28 | -0.14 | 0.441 | 0.54 |

**Table S9.** WGCNA module preservation statistics for the Leaf Gradient topological control set. Modules sorted by FDR from the genotype label permutation test used in Fig 4B (1000 shuffles of the CTRL/*Inv4m* labels). Δ*k*_*Within*_ is the mean intramodular connectivity change (*Inv4m −*CTRL); *Z*_Δ*k*_ is that observed value expressed as a *z*-score relative to the permutation null. *Z*_*density*_, *Z*_*connectivity*_, and *Z*_*summary*_ from `modulePreservation()` with 1000 permutations (Langfelder *et al*. 2011): *Z*_*summary*_ *<* 2 = not preserved, 2–10 = moderate, *>* 10 = strong. *Z*_*connectivity*_ quantifies rank preservation of intramodular connectivity; *Z*_*density*_ quantifies preservation of average connectivity strength. LFP = Largest Fragment Proportion (bootstrap support in CTRL). The Leaf Gradient set groups genes by expression along the leaf developmental gradient and serves as an orthogonal control for the Genotype Response modules in Table S8.

| Module | Genes | $Z_{density}$ | $Z_{connectivity}$ | $Z_{summary}$ | $\Delta k_{Within}$ | $Z_{\Delta k}$ | FDR | LFP |
| --- | --- | --- | --- | --- | --- | --- | --- | --- |
| coral | 26 | 3.8 | 4.8 | 4.3 | -2.35 | -1.22 | 0.220 | 0.95 |
| jade | 30 | 2.5 | 6.2 | 4.4 | -4.39 | -1.14 | 0.246 | 0.95 |
| sky | 78 | 5.1 | 10.7 | 7.9 | -13.82 | -1.09 | 0.262 | 0.83 |
| lavender | 13 | 3.6 | 4.5 | 4.1 | -1.36 | -0.72 | 0.340 | 0.81 |
| peach | 64 | 8.4 | 8.3 | 8.3 | -2.09 | -0.40 | 0.383 | 0.86 |

**Table S10.** Greenyellow module DEGs containing growth and cell division genes. Genes assigned to the greenyellow WGCNA consensus module, which has the highest bootstrap support (LFP = 0.80) among all modules. Contains the plant height GWAS candidate *sec6* and the cell proliferation marker *pcna2*. Sorted by statistical significance (FDR). GO enrichment includes “developmental cell growth” (*p* = 8.5 *×* 10^*−*4^) and “regulation of DNA replication” (*p* = 0.019).

| Gene ID | Label | Description | log<sub>2</sub>FC | FDR |
| --- | --- | --- | --- | --- |
| Zm00001eb194380 | *sec6* | Exocyst complex component Sec6 | +2.87 | $2.9 \times 10^{-17}$ |
| Zm00001eb188240 | *ropgef14* | Rop guanine nucleotide exchange factor 14 | -2.98 | $2.6 \times 10^{-14}$ |
| Zm00001eb112470 | *engd2* | OBG-type G domain containing protein | +6.14 | $4.9 \times 10^{-13}$ |
| Zm00001eb191260 | *pbl19* | Serine/threonine protein kinase PBL19 | -1.39 | $3.5 \times 10^{-9}$ |
| Zm00001eb195290 | *pap1* | PAP-associated domain containing protein | +1.45 | $4.7 \times 10^{-8}$ |
| Zm00001eb386670 | *rglg2* | E3 ubiquitin protein ligase RGLG2 | -1.01 | $5.2 \times 10^{-4}$ |
| Zm00001eb187450 | *mybr58* | MYB-related transcription factor 58 | +1.74 | $3.1 \times 10^{-3}$ |
| Zm00001eb192050 | *pcna2* | Proliferating cell nuclear antigen 2 | +2.99 | $9.3 \times 10^{-3}$ |
| Zm00001eb290800 | *pk8* | Protein kinase domain containing | -0.37 | $2.5 \times 10^{-2}$ |
| Zm00001eb392440 | *iqm1* | IQ calmodulin binding motif protein | -0.85 | $2.6 \times 10^{-2}$ |
| Zm00001eb080160 | *hipp16* | Heavy metal-associated protein 16 | -0.99 | $3.4 \times 10^{-2}$ |
| Zm00001eb404820 | *esk1* | Protein ESKIMO-like | -0.65 | $3.4 \times 10^{-2}$ |
| Zm00001eb064190 | *parg* | Poly(ADP-ribose) glycohydrolase | -0.29 | $4.2 \times 10^{-2}$ |
| Zm00001eb190580 | *pla2* | Phospholipase A2 homolog 1 | -0.69 | $4.8 \times 10^{-2}$ |
| Zm00001eb375460 | *pk3* | Protein kinase domain containing | -0.74 | $4.8 \times 10^{-2}$ |
| Zm00001eb279370 | *phi1b* | Phi-1-like phosphate induced protein | +0.72 | $5.0 \times 10^{-2}$ |

**Table S11.**
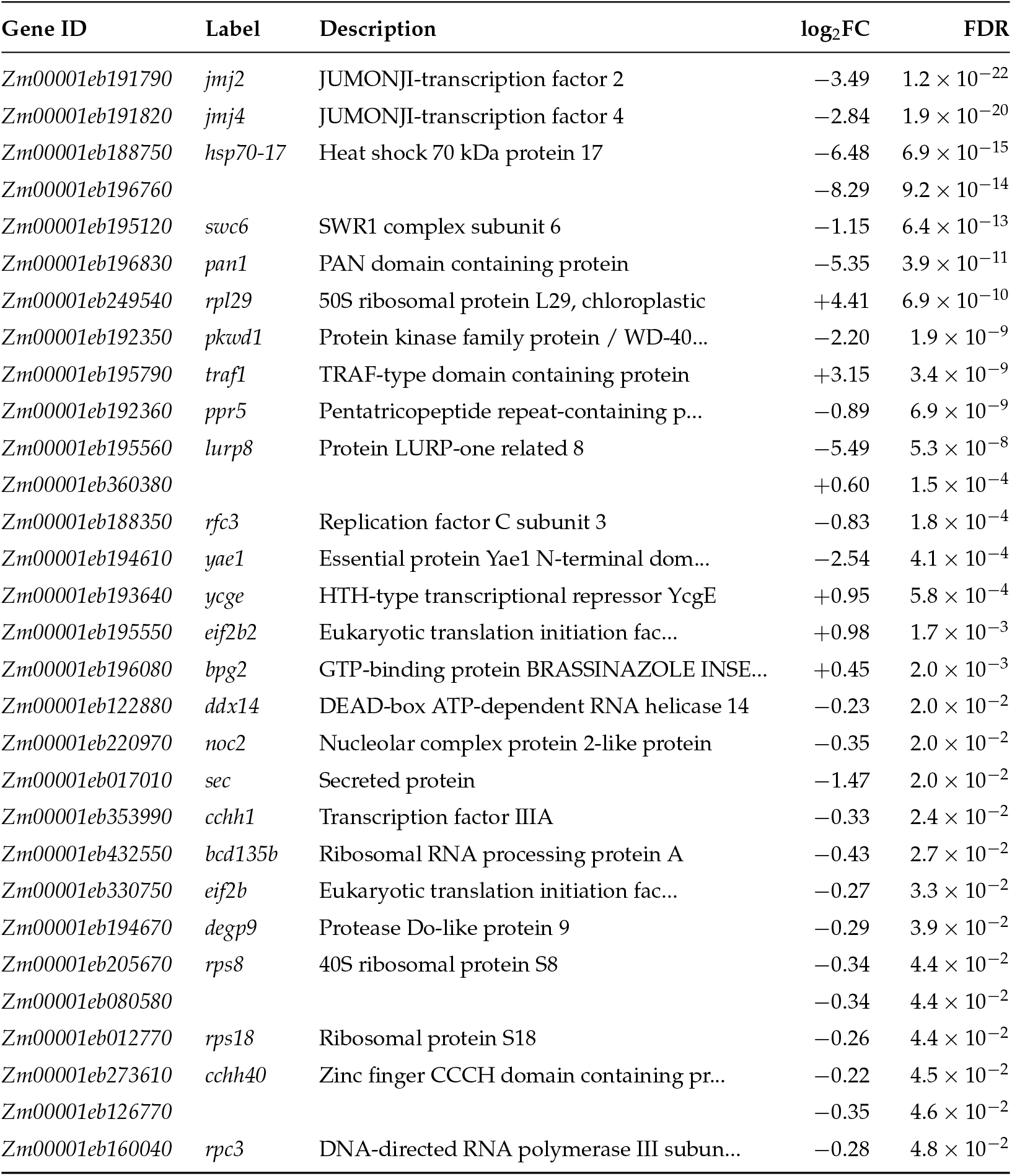
Pink module DEGs coexpressed with *jmj2*/*jmj4*. Genes assigned to the pink WGCNA consensus module, which contains the JMJ2/JMJ4 histone demethylase cluster. Sorted by statistical significance (FDR). GO enrichment for this module includes “positive regulation of growth” (*p* = 0.0017) and “ribosome biogenesis” (*p* = 0.0003).

**Table S12.**
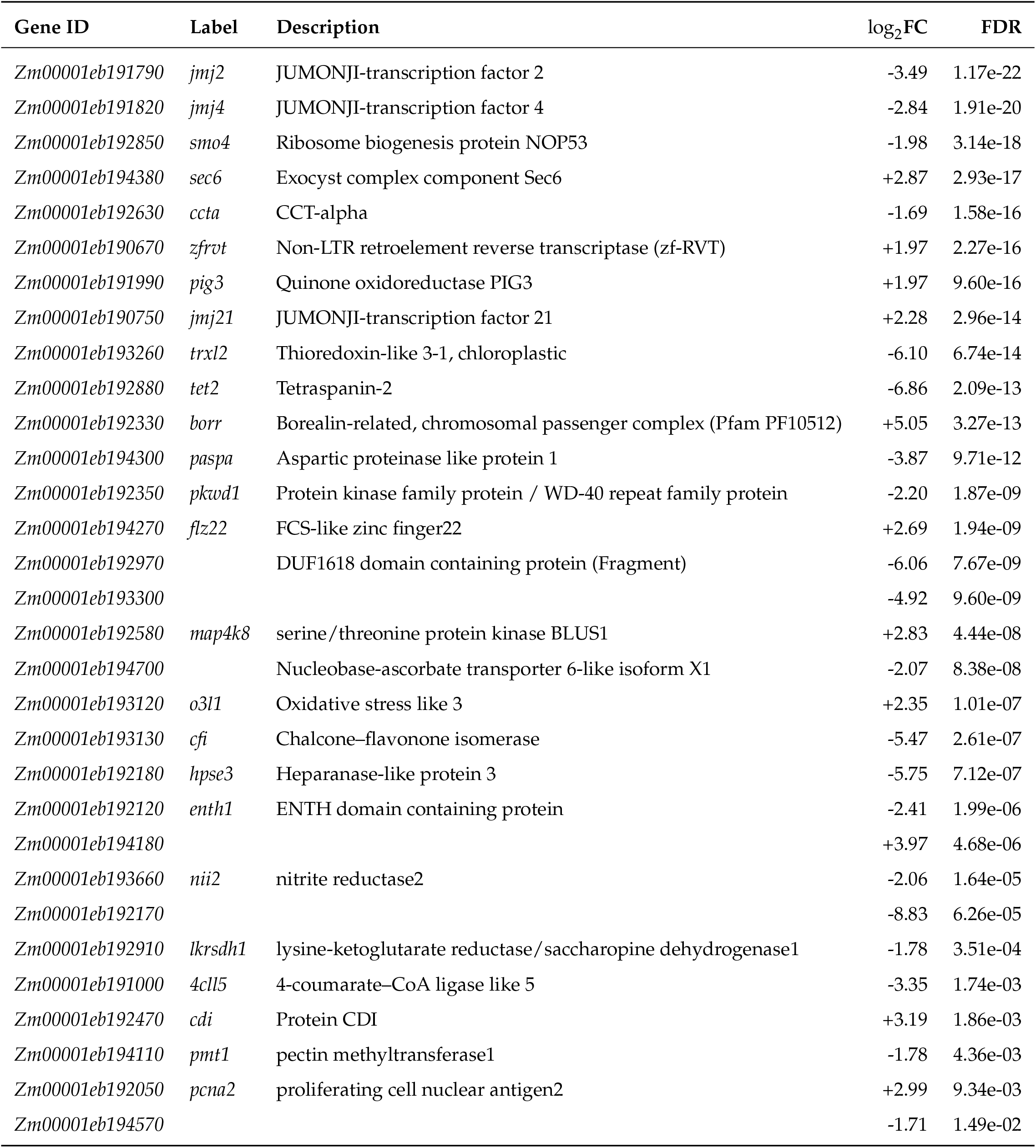
*Inv4m* trans coexpression network genes: *Inv4m* region. Genes within the *Inv4m* introgression participating in the trans coexpression network (Bonferroni *<* 0.001), spanning both MaizeNetome-supported reference edges and dataset specific novel edges, sorted by FDR ascending. log_2_FC and FDR are from the genotype DEG analysis.

**Table S13.** *Inv4m* trans coexpression network genes: outside introgression. Genes outside the *Inv4m* introgression participating in the trans coexpression network (Bonferroni *<* 0.001), spanning both MaizeNetome-supported reference edges and dataset specific novel edges, sorted by FDR ascending. log_2_FC and FDR are from the genotype DEG analysis.

| Gene ID | Label | Description | log<sub>2</sub>FC | FDR |
| --- | --- | --- | --- | --- |
| *Zm00001eb041890* | *kan1* | KANADI1 | +1.68 | 2.59e-14 |
| *Zm00001eb182850* | *mybsant* |  | +6.96 | 3.80e-13 |
| *Zm00001eb112470* | *engd2* | OBG-type G domain containing protein | +6.14 | 4.90e-13 |
| *Zm00001eb000540* | *prd4* | Protein PAIR1 (putative recombination initiation defect4) | +2.32 | 3.99e-10 |
| *Zm00001eb025970* | *pp1* | Serine/threonine protein phosphatase | +5.24 | 1.87e-09 |
| *Zm00001eb251960* | *abh1* | abscisic acid 8'-hydroxylase1 | -2.83 | 5.23e-07 |
| *Zm00001eb096620* | *pip2i* | plasma membrane intrinsic protein, PIP2i | +1.59 | 1.03e-06 |
| *Zm00001eb428740* | *ocl2* | outer cell layer2 | -2.66 | 3.43e-06 |
| *Zm00001eb157030* |  |  | +5.68 | 1.46e-05 |
| *Zm00001eb332450* | *roc4* | Peptidyl-prolyl cis-trans isomerase | -2.92 | 1.69e-05 |
| *Zm00001eb166340* | *bgal1* | Beta-galactosidase | -4.14 | 5.61e-05 |
| *Zm00001eb148670* | *acd6* | accelerated cell death ortholog6 | -1.86 | 1.51e-03 |
| *Zm00001eb183030* | *gst* | Glutathione S-transferase | +7.84 | 2.04e-03 |
| *Zm00001eb132370* |  |  | +2.33 | 4.82e-03 |
| *Zm00001eb213980* | *dyw1* | DYW family nucleic acid deaminase | -5.21 | 9.70e-03 |
| *Zm00001eb159210* | *cyp72a* | Cytochrome P450 family 72 subfamily A polypeptide 8 | -1.57 | 1.47e-02 |

**Table S14.** Full-length (Iso-seq) transcript evidence for the five B73 *jmj2*/*jmj4* cluster paralogs. Reanalysis of the Wang et al. 2020 maize Iso-seq atlas (collapsed transcripts mapped on B73 RefGen_v4) (Wang *et al*. 2020). Each v5 candidate is mapped to its v4 gene model and to the collapsed full-length, non-concatemer (FLNC) group (PB.X); columns give the FLNC read count in the three B73 inbred samples (embryo, endosperm, root) and the number of collapsed isoforms in the group. The four PB.X groups are spatially non-overlapping along the cluster, consistent with at least four independent transcription units in B73 rather than alternative isoforms of a single v5 locus. Iso-seq tissues are embryo, endosperm, and root; leaf and SAM are not sampled.

| Candidate | transcript | v4 gene | FLNC group | Isoforms | Embryo | Endosperm | Root | Total |
| --- | --- | --- | --- | --- | --- | --- | --- | --- |
| *jmj9* | Zm00001eb191790_T001 | Zm00001d051959 | PB.11785 | 6 | 6 | 9 | 6 | 21 |
| *ψ* | Zm00001d051961_T002 | Zm00001d051961 | PB.11786 | 2 | 0 | 0 | 3 | 3 |
| *jmj6* | Zm00001eb191790_T006 | Zm00001d051963 | PB.11787<sup>†</sup> | 18 | 16 | 30 | 4 | 50 |
| *jmj2* | Zm00001eb191790_T013 | Zm00001d051964 | PB.11787<sup>†</sup> | shared | — | — | — | shared |
| *jmj4* | Zm00001eb191820_T001 | Zm00001d051965 | PB.11788 | 7 | 0 | 1 | 0 | 1 |

<sup>†</sup>PB.11787 (a ~43 kb collapsed span) encompasses both *jmj6* (Zm00001d051963) and *jmj2* (Zm00001d051964), ~24 kb apart. The collapsed annotation cannot distinguish fuzzy-collapse of two separate units, a template-switch chimera, or a single long transcription unit; *jmj6* and *jmj2* counts are pooled and the *jmj2* row blanked to avoid double counting. This affects only the upper bound (four versus five paralog-level units), not the falsifier ( ≥ two). *ψ* is transcribed (three root FLNC reads) but its transcript carries a genome-encoded frameshift premature stop (B73 v5 chr4:177,445,822–177,445,824) that truncates the JMJ open reading frame, consistent with a pseudogene.

